# Exact centriole counts are critical for B cell development but not function

**DOI:** 10.1101/2024.02.14.580240

**Authors:** Marina A. Schapfl, Gina M. LoMastro, Vincent Z. Braun, Maretoshi Hirai, Michelle S. Levine, Eva Kiermaier, Verena Labi, Andrew J. Holland, Andreas Villunger

## Abstract

Centrioles define centrosome structure and function. Deregulation of centriole numbers can cause developmental defects and foster malignant disease. The p53 tumor suppressor limits the growth of cells lacking or harboring additional centrioles and can be engaged by the “mitotic surveillance” or the “PIDDosome pathway”, respectively. Here, we show that early B cell progenitors frequently present extra centrioles that are rapidly lost during maturation. Increasing centriole counts beyond physiological levels by Polo-like kinase 4 (PLK4) overexpression induces apoptosis, suggesting clearance of such cells during development. Remarkably, this apoptotic response is independent of PIDD1 or p53, but can be blocked by excess BCL2. In contrast, loss of centrosomes upon *Plk4* deletion arrests B cell development at the pro B cell stage. This defect can be rescued by co-deletion of *Usp28*, a critical component of the mitotic surveillance pathway that restores cell number and function in the absence of centrioles. In both scenarios, too many and too few centrosomes, mitochondrial apoptosis is engaged to kill B cells with abnormal centriole counts during their development with progenitor B cells being intolerant to centriole loss but permissive to centriole amplification. Unexpectedly, our findings show that centrioles are dispensable for mounting an effective humoral immune response.

## INTRODUCTION

The centrosome acts as the main microtubule-organizing center in animal cells and consists of a mature mother centriole with distal and subdistal appendages and an orthogonally attached daughter centriole, surrounded by proteinaceous pericentriolar material, the PCM (Gönczy & Hatzopoulos, 2019; Nigg & Holland, 2018). In dividing cells, the two centrioles are duplicated once during S-phase to facilitate the formation of a bi-polar mitotic spindle and thus aid the correct segregation of chromosomes. In differentiating cells, they are essential for ciliogenesis and defects in centriole biogenesis are linked to severe developmental abnormalities, including microcephaly and kidney malfunction (Mansour *et al*, 2021; Phan & Holland, 2021). Centrosome aberrations frequently occur in cancer and are known to promote chromosomal instability (CIN), provoking aneuploidy (Chunduri & Storchová, 2019; Ganem *et al*, 2009; Levine & Holland, 2018). Moreover, extra centrosomes have been linked to increased invasiveness and cancer metastasis (Arnandis *et al*, 2018; Godinho *et al*, 2014; LoMastro & Holland, 2019). As such, their number needs to be tightly regulated. Although cell division can proceed in the absence of centrioles in some circumstances, centrosomes are generally required for sustained proliferation in mammalian cells. Loss of centrioles causes delays in mitosis that activate the mitotic surveillance (*aka* stop watch) pathway, which promotes p53 stabilization by engaging the p53 binding protein, 53BP1, and the Ubiquitin Specific Peptidase, USP28. In model cell lines, USP28 activity antagonizes MDM2-mediated ubiquitination of p53, leading to its stabilization, transcription of the CDK inhibitor *p21* and cell cycle arrest in the next G1 phase (Fong *et al*, 2016; Lambrus *et al*, 2016; Meitinger *et al*, 2016). *In vivo*, mouse embryos lacking centrioles by loss of SAS4 undergo widespread p53-dependent apoptosis, around day E9.5 (Xiao *et al*, 2021). Similarly, different mutations leading to centriole biogenesis defects cause apoptosis of neural progenitor cells and microcephaly (Phan 2021 *et al, 2021*). In both scenarios, p53-induced apoptosis can be prevented by USP28 co-depletion, ameliorating the related phenotypes (Bazzi & Anderson, 2014; Phan *et al*, 2021). How p53 triggers cell death in these settings awaits detailed analysis.

Extra centrosomes, on the other hand, can engage the PIDDosome pathway, e.g., in response to cytokinesis failure or centriole amplification caused by PLK4 overexpression, leading to caspase-2-mediated cleavage of MDM2, p53 stabilization and upregulation of *p21* (Fava *et al*, 2017; Weiler *et al*, 2022). PIDDosome activation requires the interaction of PIDD1, its central component, with the transient distal appendage protein, Ankyrin Repeat Domain containing protein 26 (ANKRD26). present at the mature mother (Burigotto *et al*, 2021; Evans *et al*, 2021). PIDD1/ANKRD26 interaction is also key to control natural polyploidization of developing and regenerating hepatocytes, defining a clear signaling cascade that connects extra centrosomes to the p53 network (Sladky *et al*, 2021; Sladky *et al*, 2020). Of note, while caspase-2 has been broadly discussed to contribute to p53-induced cell death after DNA damage (Brown-Suedel & Bouchier-Hayes, 2020; Fava *et al*, 2012; Lim *et al*, 2021), apoptosis initiation in response to extra centrosomes has not been documented.

In hematopoietic cells, centrosomes exert functions that go beyond mitotic spindle pole formation and include erythroblast enucleation (Tátrai & Gergely, 2022), control of asymmetric cell division in lymphocytes (Barnett *et al*, 2012; Liedmann *et al*, 2022), immunological synapse formation (Dieckmann *et al*, 2016), as well as cell dendritic cell (DC) migration (Weier *et al*, 2022). Moreover, extra centrosomes have been documented during terminal differentiation of DCs to enhance effective helper T cell activation (Weier *et al*., 2022), and in microglia, extra centrosomes increase their efferocytosis rates (Möller *et al*, 2022). Common to these cell types is their terminally differentiated state that allows them to host extra centrosomes without endangering genome integrity, a situation similar to that found in hepatocytes (Sladky *et al*, 2022) or osteoclasts (Philip *et al*, 2022).

Whether extra centrosomes are limited to terminally differentiated innate immune cells or if they are also found in cells of the adaptive immune system has not been investigated. Hence, we set out to catalogue centriole numbers in lymphocytes along different developmental stages in primary and secondary lymphatic organs. B cells develop in the bone marrow, where they progress through defined stages that are clearly separated into phases of proliferation and differentiation (Kuppers & Dalla-Favera, 2001; Papaemmanuil *et al*, 2014; Zhang *et al*, 2011). The pro B cell and large pre B cell stage mark two early differentation stages, which are defined by fast proliferation and recombination of the immunoglobulin heavy chain *(Igh)* of the immature B cell receptor (pre BCR). Expression of the pre BCR allows cell survival and clonal expansion. Subsequently, the cell cycle is stalled, immunoglobulin light chain recombination is initiated in small, resting pre B cells to complete assembly of a mature BCR (Kuppers & Dalla-Favera, 2001; Papaemmanuil *et al*., 2014; Zhang *et al*., 2011). The successfully rearranged BCR is then expressed as cell surface-bound immunoglobuline (Ig) type M (IgM) on immature B cells that migrate to the spleen to complete initial maturation as follicular or marginal zone B cells. At this point, B cells reach competence to launch humoral immune responses in response to infection via terminal differentiation into Ig-secreting plasmablasts or plasma cells (Kuppers & Dalla-Favera, 2001; Papaemmanuil *et al*., 2014; Zhang *et al*., 2011).

We noted that proliferating B cell progenitors (pro B and large pre B cells) frequently present with additional centrioles during their ontogeny. However, those are no longer found in more mature developmental stages, beginning from the small resting pre B cell stage onwards. Apoptosis in proliferating progenitor B cells with additional centrioles can be blocked by BCL2 overexpression, suggesting clearance by apoptosis. Surprisingly, this cell death does not require p53 or PIDD1. In contrast, loss of centrioles also promotes BCL2 regulated apoptosis, yet in a strictly p53-dependent manner, arresting early B cell development. Intriguingly, B cell development and humoral immunity can occur in the absence of centrosomes as both can be restored by co-deletion of USP28. Our findings suggest that precise control of centriole counts is a prerequisite for B cell development, but not needed for their effector function in response to antigen-challenge.

## RESULTS

### Cycling B cell progenitors frequently display aberrant centriole counts

First, we monitored expression levels of key kinases involved in the centrosome cycle, PLK1, PLK2 and PLK4 (Fig. 1A) and catalogued the number of centrioles in sorted immune cell subsets along lymphocyte ontogeny by immunofluorescence using antibodies against the centriolar marker protein, CP110, and ɣ-Tubulin to localize centrosomes (Fig. S1A). Using this method, more than four centrioles were rarely found in hematopoietic stem cells (HSC), common lymphoid progenitors (CLP), developing thymocytes, as well as mature T and B cells. Remarkably, however, pro and large pre B cell progenitors showed frequent centrosome amplification with up to 25% of cycling large pre B cells displaying >4 centrioles (Fig. 1B). Of note, this phenomenon was no longer found in resting small pre B cells that exit the cell cycle to rearrange their immunoglobulin light chain locus to express a functional BCR. Signaling via the BCR is needed for maturation and survival, as well as positive and negative B cell selection processes avoiding auto-reactivity. Neither immature IgM^+^ B cells in the bone marrow nor transitional (T), mature follicular (FO) or marginal zone (MZ) B cells in the spleen, or innate-like B1 B cells in the peritoneum, displayed extra centrioles (Fig. 1B). Taking advantage of double-transgenic mice expressing the FUCCI reporter encoding geminin-GFP and Cdt1-RFP in all hematopoietic cells (Sakaue-Sawano *et al*, 2008), we could correlate increased centriole number with the mitotic activity of developing B lymphocyte progenitors where large pre B cells also presented the highest proliferation rate (Fig.1C, S1B,C). Co-staining of large pre B cells with CEP164, a marker that defines mother centrioles, suggested that the extra centrioles detected were not yet fully mature (Fig. 1D). This finding is indicative of increased nucleation activity that correlated well with higher mRNA levels of PLK1 and PLK4 at this developmental stage (Fig. 1A).

**Fig. 1.**
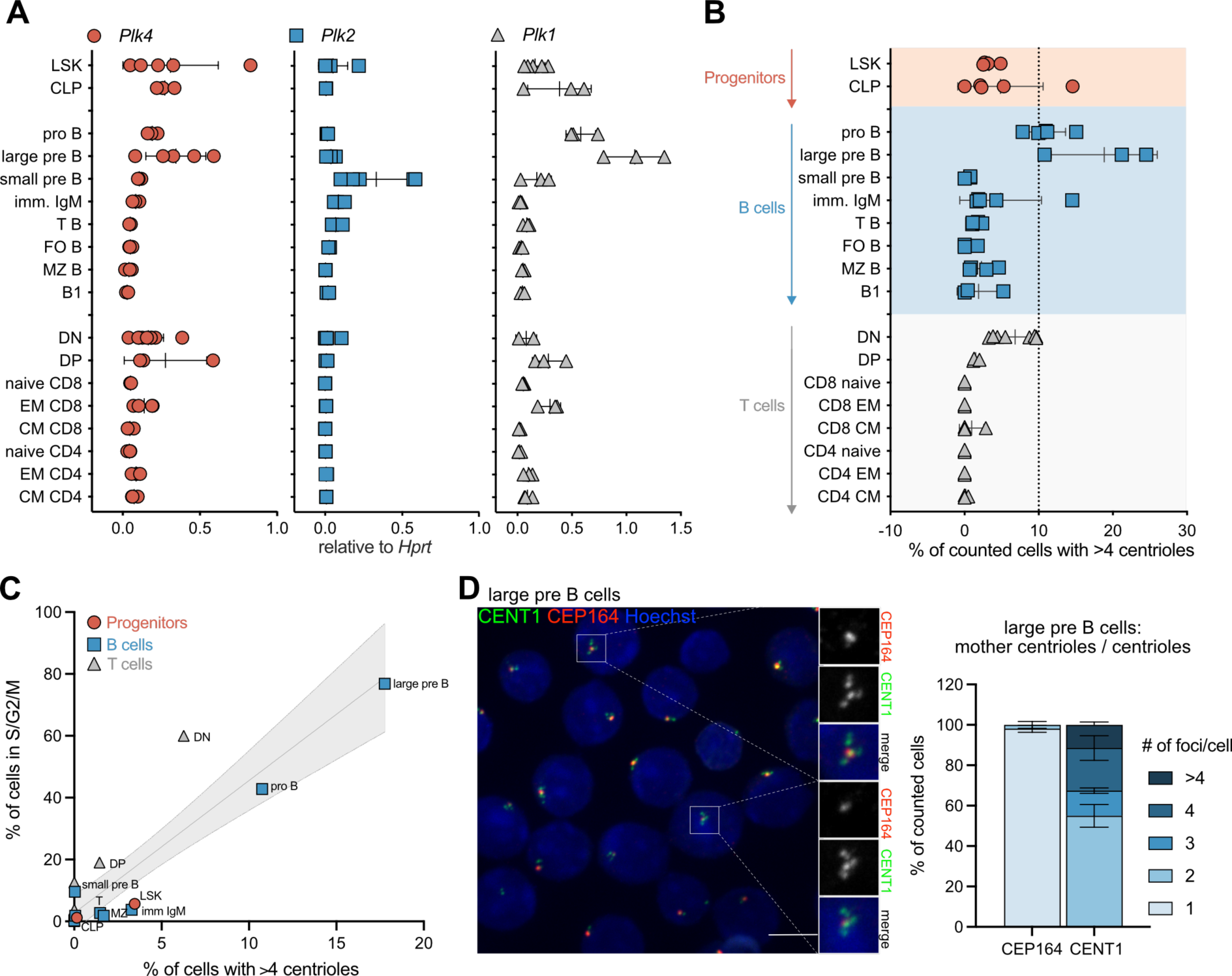
B cell progenitors display aberrant centriole numbers. A. qRT-PCR analysis for *Plk4, Plk2* and *Plk1* mRNA expression in FACS-sorted cells of the bone marrow: Lin-Sca1+c-Kit+ (LSK), common lymphoid progenitor (CLP), pro B, large and small pre B and immature IgM+ B cells; spleen: transitional (T B) B cells, mature follicular (FO B) B cells, marginal zone (MZ B) B cells, CD4 and CD8 naive, effector memory (EM) and central memory (CM) T cells; thymus: double negative (DN) and double positive (DP) thymocytes; and the peritoneum: B1 B cells (B1). B. Percentage of counted cells with >4 centrioles. Centriole number was determined by fluorescence microscopy with ɣ-Tubulin, CP110 antibody staining and Hoechst nuclear staining. C. Correlation of fraction of cells with >4 centrioles with fraction of cells in S or G2/M phase of cell cycle (SG2M), which was determined by flow cytometric analysis of mice expressing a FUCCI reporter system. Data are shown as mean of 3 individual mice. D. Immunofluorescence images and quantification of FACS-sorted large pre B cells, expressing CENT1-GFP positive daugther centrioles and stained mother centrioles with CEP164. Scale bar, 5µm. Data are shown as mean ± SD; n=3-6.

### Extra centrioles correlate with apoptosis induction in progenitor B cells

We rationalized that cycling progenitor B cells with abnormal centriole number may either undergo p21-dependent cell cycle arrest or p53-induced apoptosis. Hence, we explored the behavior of progenitor B cells in relation to centrosome number *ex vivo*. Towards this end, we isolated pro B cells from mouse bone marrow of different genotypes by cell sorting and expanded them in the presence of IL-7 to prevent their differentiation. Progenitor B cells from wt, *Pidd1^-/-^, p53^-/-^, p21^-/-^,* and *vav-BCL2* transgenic (*BCL2^tg^*) mice, expressing anti-apoptotic BCL2 in all blood cells (Ogilvy *et al*, 1998), were used. Proliferation rates after cytokine stimulation *ex vivo* were monitored by flow cytometric analyses of phospho-histone H3 levels. As noted before, BCL2 overexpressing cells showed a lower mitotic index (O’Reilly *et al*, 1997), but proliferated at rates comparable to wild type cells in response to mitogenic stimulation. p53-deficient cells seemed to proliferate slightly more on day 3, but this phenomenon did not persist at later stages (Fig. 2A, S2A). DNA content analysis showed an increase in G2/M cells in the absence of *Pidd1* or *p53*, but not *p21* (Fig. S2C,D). Analysis of ɣH2AX as a DNA damage marker revealed a significant accumulation in *BCL2* transgenic cells, but not of other genotypes (Fig. 2B, S2B). Yet, this phenomenon was not reflected in increased apoptosis. In contrast, cell death was completely blocked in BCL2 transgenic cells, while those lacking p53 or PIDD1 showed only a transient survival benefit (Fig. 2C). Of note, cell death inhibition correlated well with the maintenance of high centriole numbers in BCL2 transgenic progenitor cells (Fig. 2D, S2E). Our findings suggest that loss of *Pidd1* or *p53* are less effective in preventing centriole depletion and cell death, compared to BCL2 overexpression (Fig. 2C,D).

**Fig. 2.**
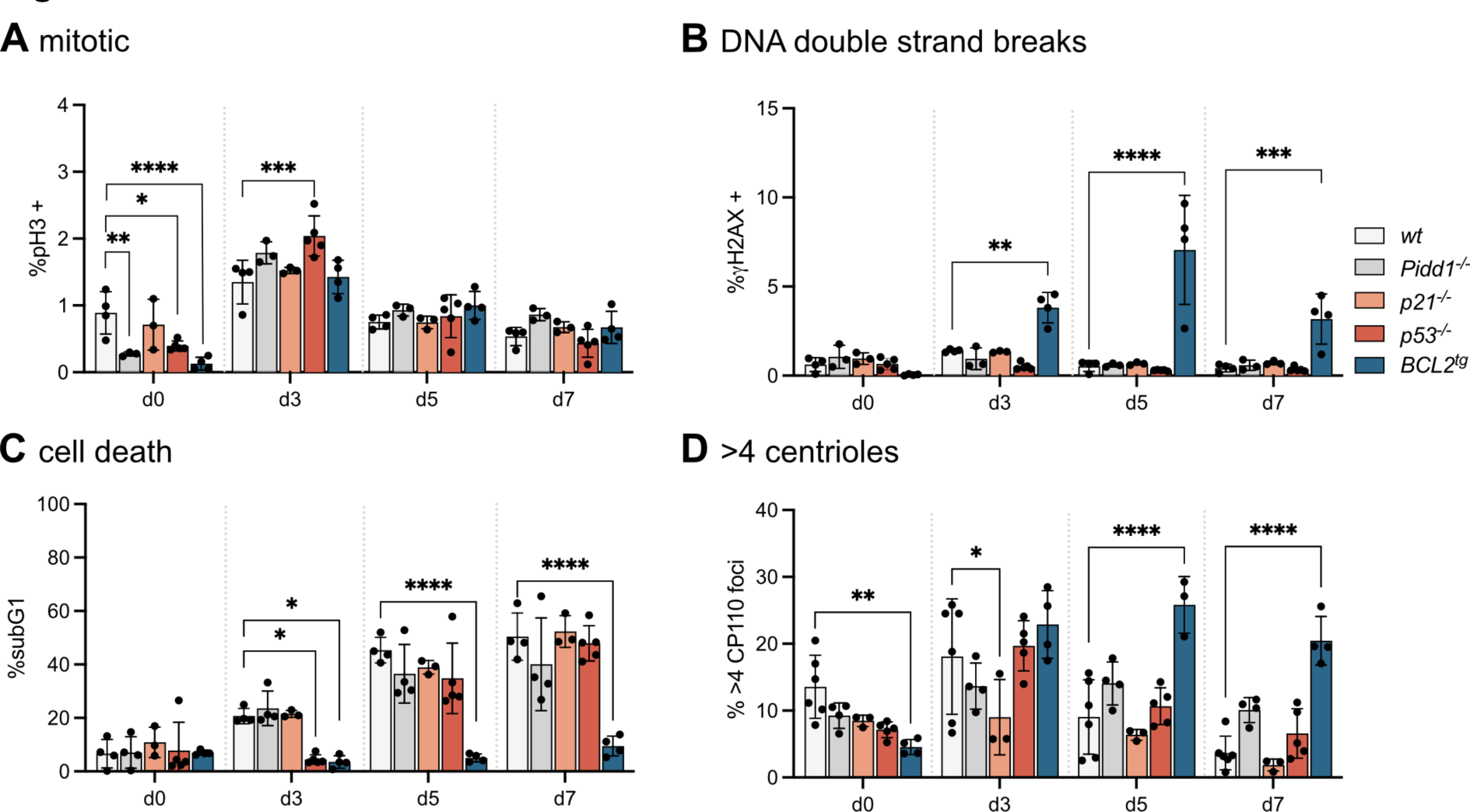
Extra centrosomes correlate with apoptosis induction in progenitor B cells. A-C. Pro B cells were FACS-sorted from mice of the indicated genotypes and flow cytometry was used to determine the fraction of (A) mitotic (pH3+) cells, (B) cells with DNA double strand breaks (ɣH2AX+) and (C) of dead cells (subG1) after sorting (d0) or in culture after 3, 5 and 7 days (d3, d5, d7) with IL7. D. Percentage of counted cells with more than 4 centrioles, determined by immunofluorescence with ɣ-Tubulin and CP110 antibody staining and Hoechst nuclear staining. Data are shown as mean ± SD; wild type *wt* (n=4-6), *Pidd1^-/-^* (n=4)*, p21^-/-^* (n=3), *p53^-/-^* (n=5), *BCL2^tg^*(n=4); *p< 0.05, **p< 0.01, ***p< 0.001, ****p< 0.0001; Genotypes were compared to *wt* by Two-way-ANOVA Tukey’s multiple comparisons test.

### PLK4 overexpression promotes progenitor B cell death

To mimic the situation seen during B cell development, we tested if overexpression of PLK4, leading to centriole overduplication, can trigger cell death. Therefore, pro B cells were isolated from the bone marrow of *Plk4*-transgenic mice, allowing conditional PLK4 overexpression in response to Doxycycline, and expanded them in IL-7. After two days, doxycycline (Dox) was added to drive excessive centriole biogenesis. *Plk4* mRNA was found increased on day 5 and day 7 leading to a detectable increase in centriole numbers (Fig. 3A,B, S3A). Assessment of cell cycle profiles, phospho-histone H3 levels and DNA damage, however, failed to reveal significant differences when compared to controls. In contrast, cell death rates were found to be increased over time in cells from *Plk4* transgenic mice (Fig. 3C, S3B-D). The fact that the cell cycle profiles did not differ suggested to us that apoptosis induction prevents pathological centriole accumulation and cell cycle arrest (Fig. S3B). Consistent with this idea, we noted that cell death was strongly reduced by BCL2 overexpression, which correlated well with an additional increase in centriole counts, while loss of PIDD1 failed to protect from cell death, a finding in line with the absence of additional mother centrioles (Fig. 1D), needed for PIDDosome formation (Burigotto *et al*., 2021; Evans *et al*., 2021).

**Fig. 3.**
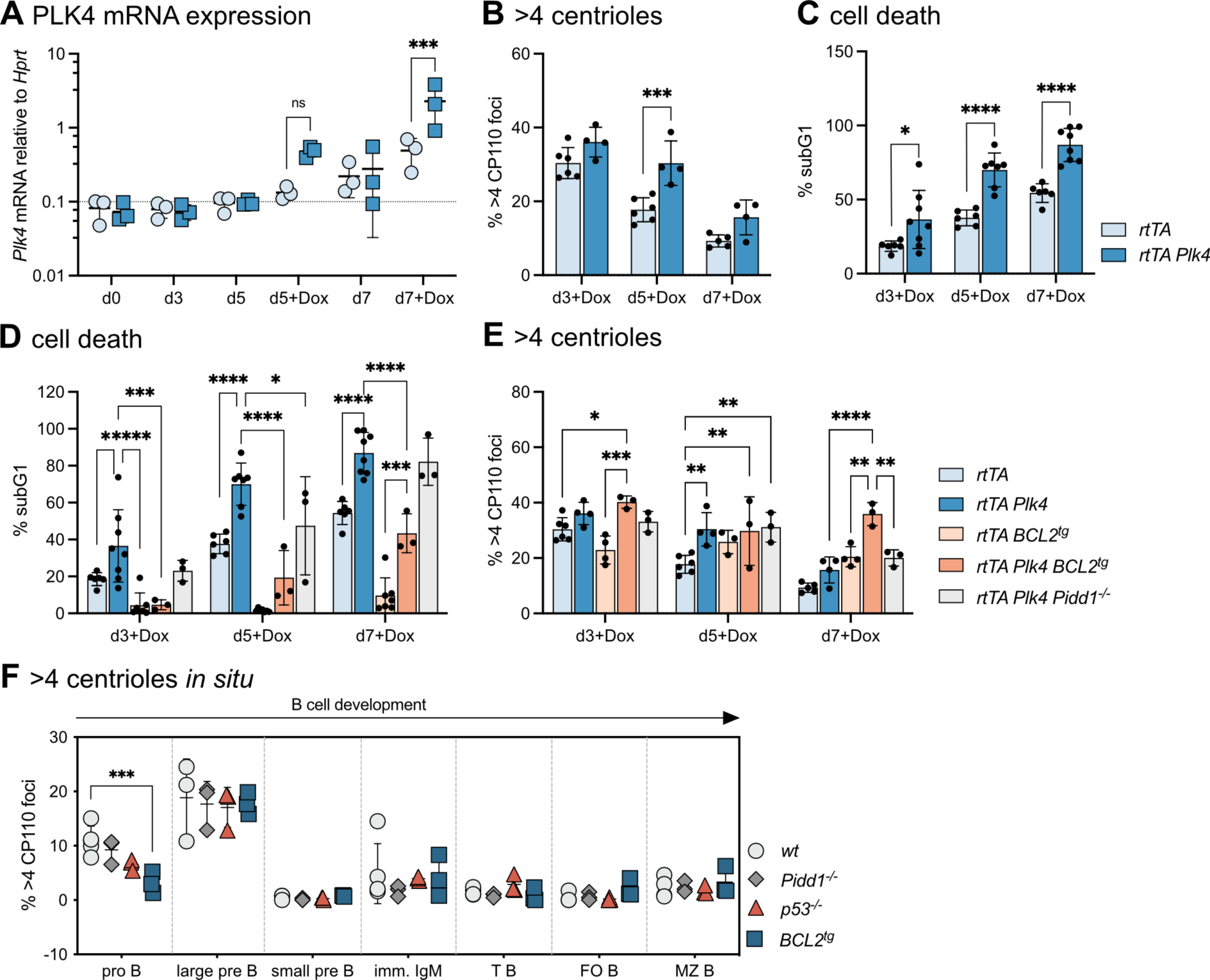
PLK4 overexpression promotes progenitor B cell death. A. Pro B cells were FACS-sorted from the indicated genotypes and put in culture. Doxycyclin was added after 48h in culture. qRT-PCR analysis of PLK4 mRNA expression in cultured pro B cells (n=3 for each genotype). B. Fraction of counted cells with more than 4 CP110 foci was determined by immunofluorescence. (n=5-6 *rtTA*, n=4 *rtTA PLK4*) C-D. Fraction of cells in in subG1 gate (cell death) was determined by flow cytometric cell cycle analysis. (n=6 *rtTA*, n=8 *rtTA Plk4*, n=7 *rtTA BCL2^tg^*, n=3 *rtTA Plk4 BCL2^tg^* and *rtTA PLK4 Pidd1^-/-^*; Genotypes were compared to *rtTA Plk4 or rtTA BCL2^tg^* for statistical analysis). E. Fraction of counted cells with more than 4 CP110 foci was determined by immunofluorescence. (n=5-6 *rtTA*, n=4 *rtTA Plk4*, n=3-4 *rtTA BCL2^tg^,* n=3 *rtTA Plk4 BCL2^tg^* and *rtTA PLK4 Pidd1^-/^*) F. Immunofluorescence with CP110, ɣ-Tubulin of FACS-sorted B cells was used to determine the fraction of cells with more than 4 centrioles of *wt* (n=4), *BCL2^tg^* (n=3) and *p53^-/-^* (n=3) animals. Bone marrow: pro B, large pre B, small pre B and immature IgM+ (imm. IgM) B cells; Spleen: transitional (T B), mature follicular (FO B) and marginal zone (MZ B) B cells. Data are shown as mean ± SD; *p<0.05, **p<0.01, ***p<0.001, ****p<0.0001; Two-way-ANOVA Tukey’s multiple comparisons test.

We wondered if we might detect extra centrioles at later developmental B cell stages *in situ* when apoptosis or cell cycle arrest were perturbed. Different B cell differentiation stages were isolated by cell sorting from wt, *Pidd1^-/-^*, *p53^-/-^*, or *Vav-BCL2* transgenic mice and analyzed for centriole numbers using CP110 and ɣ-Tubulin staining to localize centrosomes by IF. Remarkably, none of the genotypes analyzed showed centriole counts that exceeded those found in wild type progenitor B cells or later developmental stages (Fig. 3F). Taken together, our findings document BCL2-regulated cell death as a barrier against aberrant centriole numbers in progenitor B cells *ex vivo*, but additional mechanisms must prevent their accumulation *in vivo*.

### Centriole loss promotes p53-dependent apoptosis in early B cells

Additional centrioles do not accumulate along B cell development, as carriers are effectively cleared by BCL2-dependent and independent mechanisms *in vivo*. However, the impact of centriole loss on B cell development has not been assessed. Hence, we explored the response of progenitor B cells to PLK4 inhibition using centrinone (Wong *et al*, 2015). In cancer cell lines, subsequent centriole depletion causes mitotic delays due to problems with proper spindle pole formation (Levine & Holland, 2018). This activates the mitotic surveillance pathway to promote p53-induced cell cycle arrest but can also be cytotoxic to neuronal progenitor cells (Phan *et al*., 2021) or breast cancer cells displaying high TRIM37 E3 ligase activity (Meitinger *et al*, 2020; Yeow *et al*, 2020). Here, we included *p21* and *p53* mutant as well as BCL2 overexpressing cells in our analysis. Addition of centrinone to pro B cell cultures reduced centriole count in a dose and time-dependent manner in all genotypes alike (Fig. 4A,B, S4A). DNA content and phosphor-histone H3 analyses suggested that loss of *p21* or *p53* weakened the mitotic surveillance pathway, as expected (Fig. 4C, S4B), leading to an increase of the percentage of cells in mitosis, albeit *p21* loss did not yield statistical significance (Fig. S4B). BCL2 overexpressing cells were also found to show an increased mitotic index, presumably due to abrogated cell death signaling (Fig. S4B). Of note, loss of p53 reduced the number of apoptotic cells to a similar degree as BCL2 overexpression, rendering cells highly resistant to centrinone treatment and allowing also the survival of polyploid cells (Fig. 4D, S4C). Together, this suggests that p53 is the dominant route to apoptosis after centriole depletion.

**Fig. 4.**
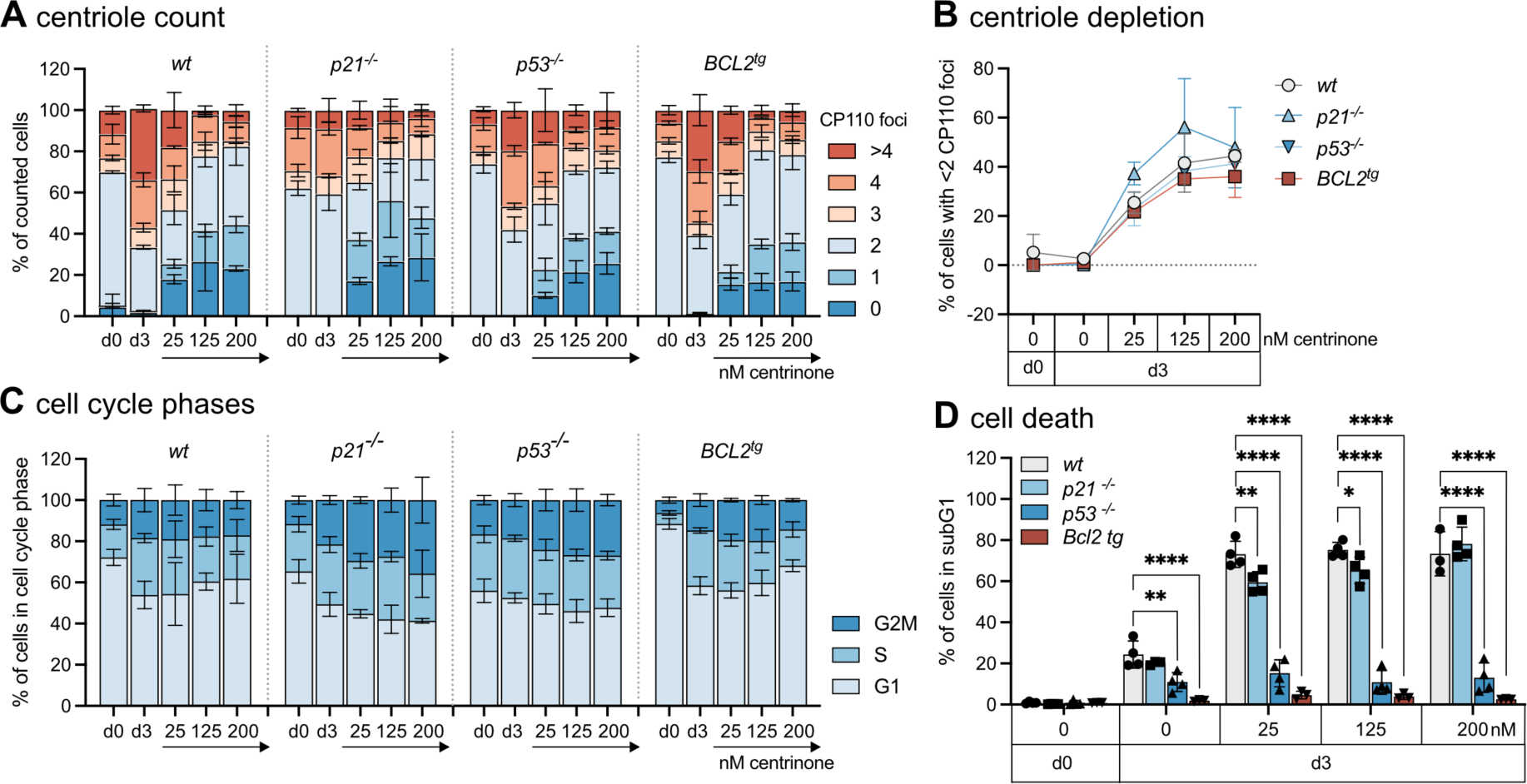
Centriole loss promotes p53-dependent apoptosis in early B cells. A. FACS-sorted pro B cells of *wt* (n=4), *p21^-/-^* (n=4), *p53^-/-^*(n=4) and *BCL2^tg^* (n=3) were treated with 25, 125 or 200nM centrinone for 3 days in presence of IL-7. Centriole distribution and (B) depletion (<2 centrioles) was assessed by immunofluorescence with ɣ-Tubulin, CP110 antibodies and Hoechst staining. C. Fraction of cells in G1-, S-, G2/M-phase of the cell cycle and (D) fraction of cells in subG1 gate (cell death) was determined by flow cytometric cell cycle analysis. Data are shown as mean ± SD; n=3-5; *p<0.05, **p<0.01, ***p<0.001, ****p<0.0001; Genotypes were compared to *wt* by Two-way-ANOVA Tukey’s multiple comparisons test.

### Loss of PLK4 arrests early B cell development

To corroborate these findings *in vivo*, we analyzed B cell development in mice expressing a floxed *Plk4* allele in combination with the *Mb1^Cre^* transgene, allowing target gene deletion in the B cell lineage. Analysis of bone marrow and spleen revealed a clear reduction in the overall percentage and number of B cells. Analysis of bone marrow revealed that loss of PLK4 expression led to a severe developmental block at the pro/pre B cell stage with a concomitant drop in immature IgM^+^D^-^ and IgM^+^D^+^ mature recirculating B cells (Fig. 5A). A more detailed analysis of progenitor B cell stages documented an accumulation at the pro B cell stage where cells are highly proliferative and the *Mb1^Cre^* allele becomes active. Within the pre B cell pool, there was a significant drop in small resting and a relative increase in large cycling pre B cells, indicating that these cells may not be able to exit the cell cycle, presumably because they may die off at this stage (Fig. 5A). Corresponding changes were also noted in absolute cell numbers (Fig. S5A) and sorted pro B cells lacking *Plk4* did not thrive in the presence of IL-7 *ex vivo* (Fig. S5B). Consistent with a differentiation defect linked to increased cell death, *Mb1^Cre^Plk4^F/F^* mice showed a significantly reduced B cell percentage and number in the spleen, with all transitional (T1, T2) and terminal differentiation stages (FO, follicular and MZ, marginal zone B cells) similarly affected (Fig. 5B, S5C).

**Fig. 5.**
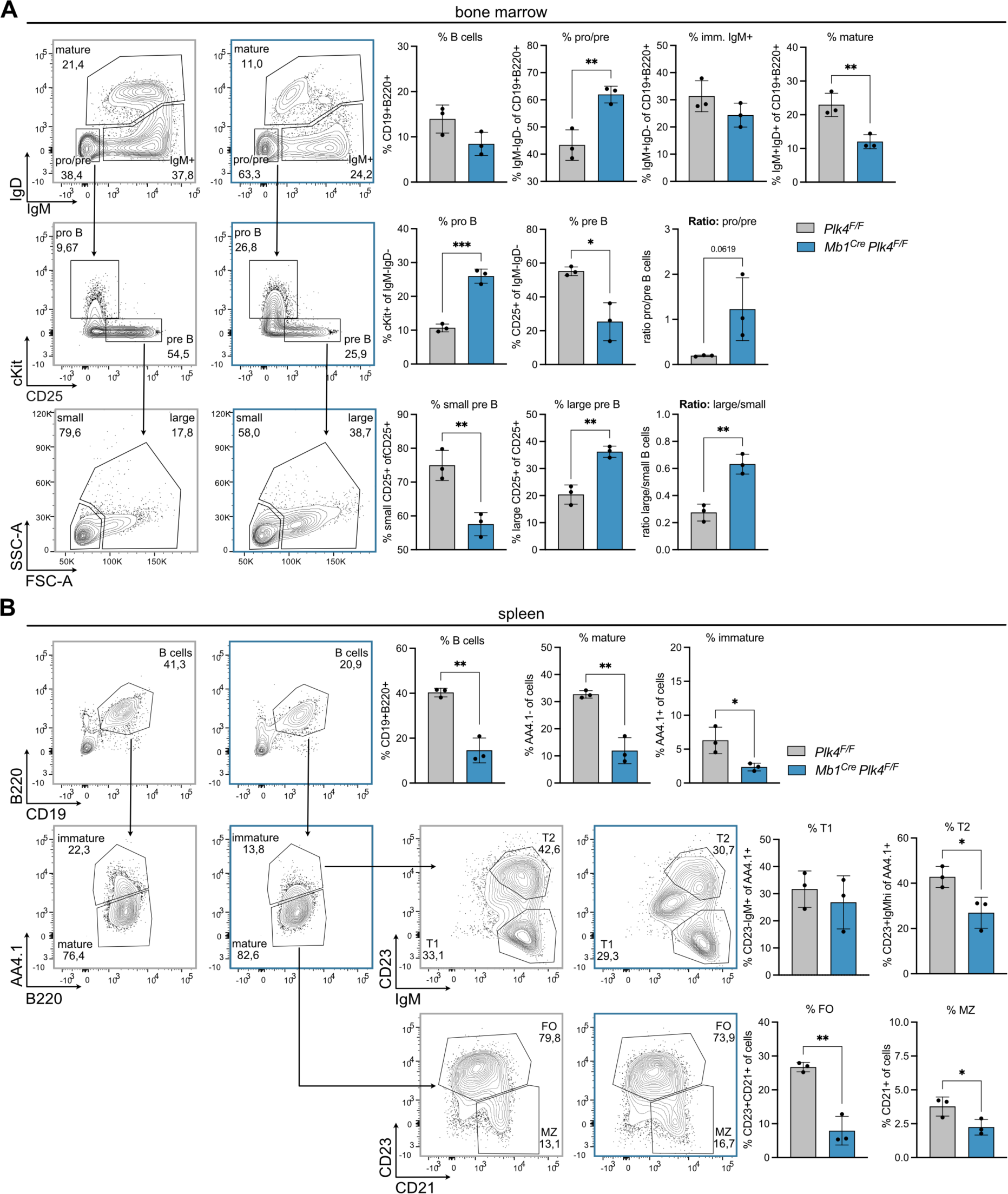
Loss of PLK4 arrests B cell development. A. Representative FACS plots illustrating the gating of pro/pre-B (IgD-IgM−),immature (IgD-IgM+) and mature B cells (IgD+IgM+) in the bone marrow after pregating on CD19+B220+ cells; from pro/pre gate further gating on pro B (cKit+CD25-) and pre B (cKit-CD25+) cells; and from pre B further gating on small and large pre B cells via FSC-A and SSC-A. Bar graphs show the fraction of B cells, pro-/pre, immature, and mature B cells within the B220+CD19+ B cell population; fraction of pro and pre and ratio of pro to pre B cells within the pro/pre B cell population and fraction of small and large pre B and ratio of large to small pre B cells within the pre B cell population of control (*Plk4^F/F^*) and *Mb1^Cre^ Plk4^F/F^* mice. B. Representative FACS plots illustrating the gating of splenic B cells (CD19+B220+); immature (AA4.1+) and mature B cells (AA4.1-); from mature B cell gate further gating on follicular (FO, CD23+) and marginal zone (MZ, CD21+CD23-); and from immature B cell gate further gating on transitional 1 (T1, IgM+CD23-) and transitional 2 (T2, IgM+CD23+) B cells. Bar graphs show the fraction of B cells, mature, immature, FO, MZ, T1 and T2 cells within the population of all cells of control (*Plk4^F/F^*) and *Mb1^Cre^ Plk4^F/F^* mice. Error bars depict the standard deviation of the mean; n=3; *p<0.05, **p<0.01, ***p<0.001, ****p<0.0001; Unpaired T-test.

### Blunting the mitotic surveillance pathway rescues B cell survival and function

Based on our prior analysis we reasoned that activation of the mitotic surveillance pathway might trigger early progenitor B cell death. Hence, we attempted a genetic rescue experiment by co-deleting USP28. Remarkably, while *Mb1^Cre^Plk4^F/F^* mice showed a near complete loss of CD19^+^ B cells in peripheral blood with a parallel increase in CD3^+^ T cells, co-deletion of USP28 restored the percentage of CD19^+^ cells (Fig. 6A). Similarly, the fraction of pro/pre B cells, CD25^+^ pre B cells and large pre B cells in the bone marrow, as well as the percentage of B cells in the spleen, were rescued to levels seen in *Plk4^F/F^Usp28^F/F^* control mice (Fig. 6B,C). Analysis of steady state serum immunoglobulin levels revealed that *Mb1^Cre^Plk4^F/F^* mice still produced near normal levels of IgM, while IgG1 levels were significantly reduced. In *Mb1^Cre^Plk4^F/F^Usp28 ^F/F^* mice, however, both IgM and IgG1 levels were normalized again. Hence, we tested whether centriole-depleted B cells were still able to mount an adaptive immune response and immunized animals with the T cell dependent model antigen, ovalbumin (OVA). Remarkably, while *Mb1^Cre^Plk4^F/F^* failed to produce OVA specific immunoglobulins and a concomitant Increase In spleen weight, *Mb1^Cre^Plk4 ^F/F^Usp28 ^F/F^* managed to do so, and reached OVA-specific Ig levels comparable to those seen in control mice (Fig. 6C).

**Fig. 6.**
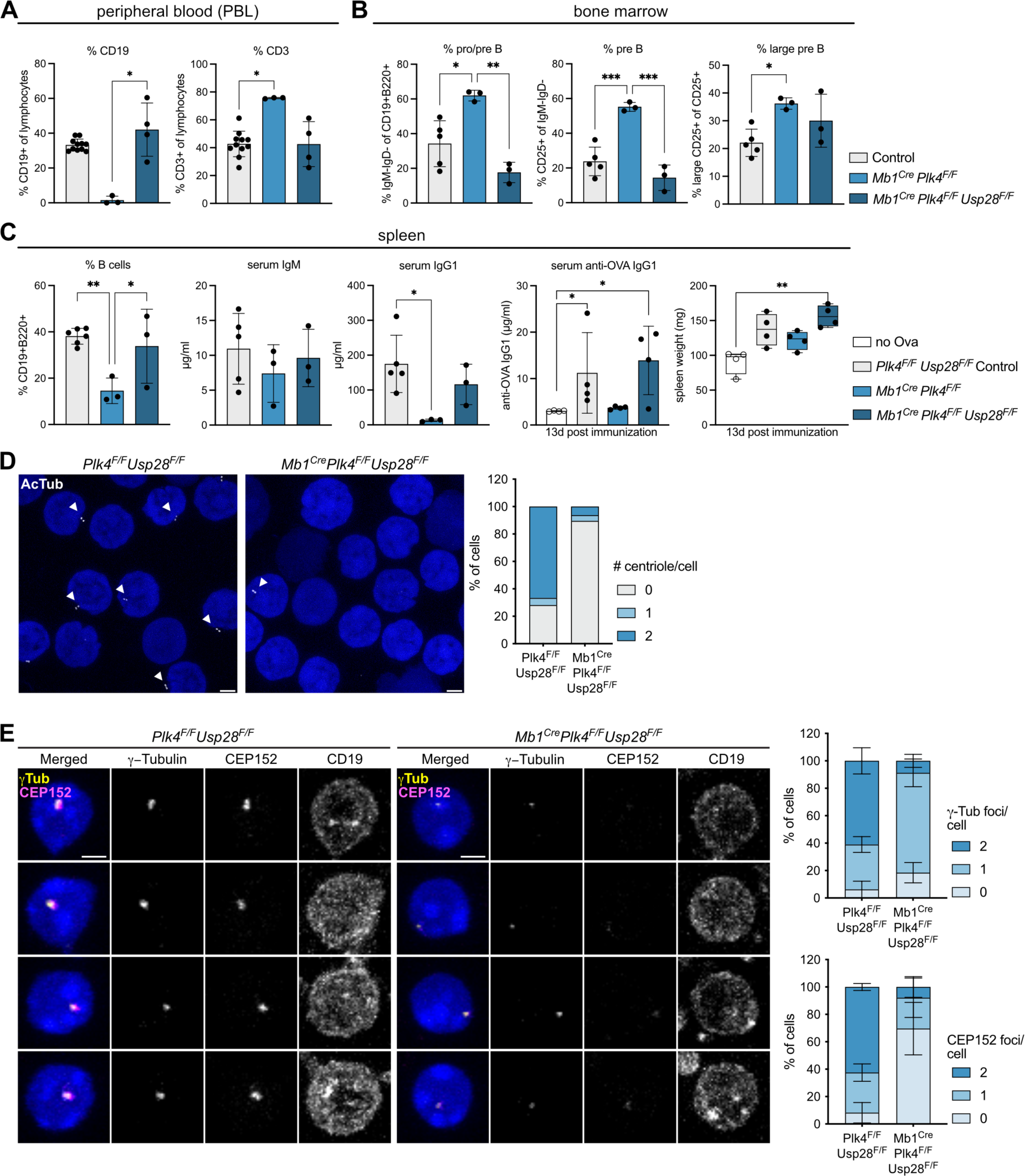
Co-depletion of USP28 restores B cell development in the absence of PLK4. A. Percentage of CD3+ or CD19+ cells in the peripheral blood of *Plk4^F/F^* or *Mb1^Cre^* control mice (n=11) and *Mb1^Cre^ Plk4^F/F^* (n=3) and *Mb1^Cre^ Plk4^F/F^ Usp28^F/F^*mice (n=4) determined by flow cytometry. B. Fraction of pro/pre B (IgM-IgD-of CD19+B220+), pre B cells (CD25+ of IgM-IgD-) and large pre B cells (FSC-A^hi^SSC-A^hi^ of CD25+) from the bone marrow of *Plk4^F/F^* or *Plk4^F/F^ Usp28^F/F^* control animals (n=5-6), *Mb1^Cre^ Plk4^F/F^* (n=3) and *Mb1^Cre^ Plk4^F/F^ Usp28^F/F^*(n=3). C. Fraction of B cells (CD19+B220+ of cells) of the spleen determined by flow cytometry and serum IgM, IgG1 levels determined by ELISA of unchallenged animals. Serum anti-OVA-IgG1 and spleen weight of indicated genotypes were analyzed 13 days post immunization (n=4 for all genotypes, as control homozygous and heterozygous *Plk4* and *Usp28* floxed mice were used). D. Expansion microscopy was used to determine the number of centrioles per cell in *PLK4^F/F^ Usp28^F/F^* and *Mb1^Cre^ Plk4^F/F^ Usp28^F/F^* mice in splenic B cells (isolated via negative selection, n=96 cells from 1 mouse). Cells were expanded by a factor of 4 and stained with acetylated Tubulin (AcTub) antibody and DAPI. Scale bars represent 2.5µm. E. Number of CEP152 and ɣ-Tubulin foci per cell was determined by immunofluorescence of CD19+ B cells of the peripheral blood isolated from *PLK4^F/F^ Usp28^F/F^* and *Mb1^Cre^ Plk4^F/F^ Usp28^F/F^* mice. Cells were stained with CD19, CEP152, ɣ-Tubulin antibodies and DAPI (representative IF images) (n=3). Data are shown as mean ± SD (except error bars for spleen weight depict min to max); *p<0.05, **p<0.01, ***p<0.001, ****p<0.0001; One-way-ANOVA Tukey’s multiple comparisons test.

To confirm that these B cells indeed can survive and mature in the absence of centrioles, we conducted IF staining experiments in CD19^+^ cells isolated from spleens of *Mb1^Cre^Plk^F/F^ Usp28^F/F^*mice. Staining for acetylated Tubulin (AcTub), ɣ-Tubulin, CEP135 or CEP152 all confirmed loss of centrioles in the majority of B cells lacking *Plk4* (Fig. 6D,E, S6A). Moreover, expansion microscopy documented frequent structural alterations in the rare remaining centrioles found in B cells from *Mb1^Cre^Plk ^F/F^Usp28 ^F/F^* mice (Fig. S6B).

Finally, we corroborated these findings in an in vitro co-culturing system of naïve B cells and CD40 ligand expressing and BAFF-secreting feeder cells, mimicking an induced germinal center (iGC) reaction (REF). This allowed us to determine the dependency of iGC B cells on centrioles for class switch recombination (CSR) and plasmablast generation. Consistent with mouse genetics, centriole loss induced by centrinone treatment did not impact cell survival, IL-21-driven differentiation into CD138+ plasmablasts or CSR from IgM to IgG1 of GC B cells (Fig. S6C-E).

While we could confirm the loss of centrioles via immunofluorescence of GC B cells after 4 days, the cell cycle activity was comparable in the presence or absence of centrinone (Fig. S6F-G).

Together, this suggests that mature B cells can proliferate, undergo class switch recombination and differentiate into plasmablast to secrete immunoglobulins upon antigenic challenge in the absence of centrioles.

## DISCUSSION

Here, we provide evidence that B cell development depends on the maintenance of exact centriole counts and that amplified centrioles are a regular feature of cycling pro and large pre B cells. Additional centrioles, however, are no longer detectable at more mature B cell differentiation stages and subsets, starting from small resting pre B cells onwards (Fig. 1). We believe that their clearance is critical to maintain genome integrity as their presence coincides with the differentiation stage where early B cell malignancies frequently arise (Greaves, 2018; Jackson *et al*, 2021). Consistently, centrosomal abnormalities are a recurrent feature of pre B acute lymphoblastic leukemia in humans, and correlate with poor prognosis but not with a particular genetic makeup of ALL (Guo *et al*, 2022; Kerketta *et al*, 2017). Whether extra centrioles exert a biological function at this developmental B cells stage or if they mark non-functional cells at risk to fail development or transform remains unclear. We initially reasoned that such cells may have experienced delays or errors during heavy chain rearrangement, leading to persisting DNA damage, cell cycle delays and subsequent cell death. Of note, p53 pathway activity is actually repressed at this stage epigenetically by the Polycomb group protein, BMI, to allow proliferation and *Ig* heavy chain recombination (Cantor *et al*, 2019). This may explain why neither loss of p53 itself, nor PIDD1, that can act upstream of p53 (Fava *et al*., 2017), provide prolonged cell death protection comparable to that of BCL2 overexpression in *ex vivo* cultures (Fig. 2). Moreover, while B cells overexpressing BCL2 survive long term *ex vivo* and display extra centrioles, *Rag1*-deficient progenitor B cells, unable to recombine DNA at this developmental stage, do not (Fig. S2, not shown). This argues against secondary consequences of faulty *Igh* rearrangement and physiological DNA damage as triggers. Moreover, a significant increase in DNA double strand breaks was only noted in BCL2 transgenic cells (Fig. 2), consistent with their ability to survive in their presence (Strasser *et al*, 1994). In line with our observations made in *ex vivo* cultures, excessive centriole biogenesis due to PLK4 overexpression triggered BCL2 regulated apoptosis (Fig. 3). Surprisingly, loss of *Pidd1* did not prevent this, despite PIDD1 being activated in response to PLK4 overexpression in different cell line models (Burigotto *et al*., 2021; Evans *et al*., 2021). Unresponsiveness may again be related to the fact that p53 is repressed at this developmental stage to allow expansion of progenitor B cells (Cantor *et al*., 2019). Of note, extra centrioles have been observed also in multi-ciliated olfactory sensory neurons (OSN) and their immediate neuronal precursors, that can still cycle in their presence. Amplification was attributed to high levels of the *SCL/Tal1-interrupting locus* gene (*Stil*) and *Plk4* (Ching & Stearns, 2020). *Plk1* and *Plk4* also appear most abundant in cycling progenitor B cells (Fig. 1). Whether the increased mRNA expression levels of centriole biogenesis genes noted are indeed causal for the centriole amplification seen in OSNs or if they solely correlate with the proliferative activity across different progenitor cell types remains to be investigated further. Strikingly though, extra centrioles were not observed in later differentiation stages of B cell development, neither in the absence of PIDD1, p53 nor the presence of transgenic BCL2 (Fig. 3). This suggests that alternative mechanisms are engaged to clear such cells *in vivo* during B cell ontogeny, opening new lines of investigation. At the moment, we can only speculate that other cell death types may kick in to clear such cells in the bone marrow, or that centrioles are themselves cleared, e.g. by macro-autophagy.

In contrast, loss of centrioles induced by chemical inhibition of PLK4, leading to the activation of the mitotic surveillance pathway, also triggers BCL2 regulated apoptosis *ex vivo* that is strictly p53 dependent (Fig. 4). This is surprising, given the proposed developmental repression of p53 in progenitor B cells (Cantor *et al*., 2019). However, in this scenario, mitotic delays after centrinone treatment must lead to p53 accumulation and this appears to induce transcriptional activation of target genes, promoting BCL2-regulated apoptosis. Likely candidates and p53 targets include *BAX* (Miyashita & Reed, 1995)*, PUMA/BBC3* and *NOXA/PMAIP* (Nakano & Vousden, 2001; Oda *et al*, 2000). Consistent with the detrimental effects of PLK4 inhibition on progenitor B cell survival, we noted developmental arrest of B cells at the pro B cell stage upon *Mb1^Cre^*-driven deletion of *Plk4*, correlating with increased cell death *ex vivo* and leading to a severe drop in the percentage and number of all subsequent B cell subsets in bone marrow and spleen of *Mb1^Cre^Plk4^f/f^* mice (Fig. 5, S5). This cell loss could be largely corrected by co-deletion of *Usp28*, leading to restoration of B cell numbers in bone marrow and spleen (Fig. 6). Our findings are in line with studies demonstrating the rescue of neuronal progenitor cells lost due activation of the mitotic surveillance pathway by co-deletion of USP28 or p53 (Phan et al., 2021). Loss of *Usp28* was also shown to rescue development in early mouse embryos lacking the centriolar assembly factor SAS4 (Bazzi & Anderson, 2014; Xiao *et al*., 2021). Conditional deletion of *Sas4* revealed a similar cell death response during lung development (Xie *et al*, 2021). Together, these findings place progenitor B cells on the list of cell types that critically depend on the surveillance of exact centriole counts and that respond with apoptosis upon their loss. This feature appears far from universal, as centriole deficiency does not affect developmental, neonatal or compensatory hepatocyte proliferation in mice lacking *Plk4* in the liver (Sladky *et al*., 2022), nor that of gastrointestinal stem cells in the absence of *Sas4* (Xie *et al*., 2021). Mechanisms underlying these cell type or tissue-dependent differences remain to be uncovered.

Surprisingly, loss of USP28 not only restored B cell maturation and survival, but also B cell function, as indicated by immunoglobulin production in steady state but also upon OVA-challenge (Fig. 6C). While loss of *Plk4* did not significantly affect IgM immunoglobulin levels, representing largely natural antibodies that are produced in the absence of infection, IgG1 levels clearly dropped in *Plk4* mutant mice, but were back to normal levels when USP28 was co-deleted (Fig. 6C). The latter suggest defects in class switch recombination in germinal center reactions, presumably due to the death of expanding antigen-stimulated naïve B cells or during differentiation into plasmablasts or plasma cells. This may be linked to the inability of these cells to conduct asymmetric cell division during germinal center reactions (Barnett et al., 2012). Moreover, our OVA-immunization experiment indicates that acentrosomal B cells can still undergo class switch recombination, producing comparable levels of OVA-specific IgG1, and hence must be able to expand and survive in the absence of centrosomes, when p53 signaling is blunted. Similar findings have been made in NPCs carrying mutations affecting centriole biogenesis and mitotic timing (Phan *et al*., 2021). Progenitor B cells appear to share this capacity. Given the proposed role of the centrosome in lysosome positioning for antigen-extraction at the immunological synapse in B cells (Yuseff *et al*, 2011) it was interesting to see that *Mb1^Cre^Plk4^F/F^Usp28^F/F^*mice appear to mount a normal IgG1 response to immunization (Fig. 6C). If these immunoglobulins are equal in affinity to OVA-specific antibodies produced in control animals was not assessed. Regardless, our IF analysis of CD19^+^ positive B cells from the blood documents the absence of centrosomes in the residual B cells found in the absence of *Plk4*. Notably, the few remaining centrioles also displayed structural defects (Fig. 6), and hence were unlikely fully functional. Moreover, our iGC culture analyses in the presence of centrinone support the notion mature B cells do not rely on functional centrosomes for activation, differentiation or CSR (Fig. S6). This argues against counterselection and expansion of residual B cells that may have escaped homozygous *Plk4* deletion. As such, we propose that mature B cells experiencing centriole depletion can still proliferate and expand upon antigenic challenge.

Together, our results suggest that B cells are intolerant to centriole loss but permissive to centriole accumulation. While the cause for this permissiveness and the trigger for centriole accumulation nor their potential physiological function in progenitor B cells are not yet known, it is tempting to speculate that malignant transformation of these cells, most frequently seen in early childhood, may be facilitated by the survival of cells carrying extra centrioles. Open questions to address in future studies are how extra centrioles are cleared during B cell ontogeny in vivo and if the absence of centrosomes may affect antibody affinity maturation, given their documented role in antigen extraction and presentation on B cells.

## Materials & Methods

### Animal models

Breeding colonies were approved by the Austrian Federal Ministry of Education, Science and Research (BMWF: 66.011/0008-V/3b/2019), or approved by the Johns Hopkins University Institute Animal Care and Use Committee (MO21M300). Generation and genotyping of *R26-rtTA, TET-Plk4* (Levine et al, 2017), *Centrin-GFP* (Hirai et al, 2016), *Vav-BCL2* (Ogilvy et al., 1998), *p53^-/-^* (Lowe *et al*, 1993) and *p21^-/-^*(Deng *et al*, 1995) mice were described before. These mice were backcrossed and maintained on a C57BL/6N background for at least 12 generations and housed at the animal facility of the Medical University of Innsbruck under SPF conditions. *Usp28^F/-^* mice were obtained from the laboratory M. Eilers (Diefenbacher *et al*, 2014). *Plk4^F/F^* mice were generated as described previously (LoMastro *et al*, 2022) and are available at The Jackson Laboratory Repository (Stock #037549). *Mb1^Cre^* mice were obtained from The Jackson Laboratory Repository (Stock #020505). These mice were kept on a mixed background and were housed and cared for in an AAALAC-accredited facility. For immunization, mice were injected i.p. with 100 µg Endofit Ovalbumin (Invivogen 9006-59-1) mixed in a 1:1 ratio with Imject^TM^ Alum Adjuvant (Thermo Fisher 77161) in a volume of 200µL per mouse. For all experiments male and female littermate mice were used indiscriminately, except for *p53^-/-^*animals, which were only male.

### Preparation of single-cell suspensions

For the generation of single-cell suspensions murine organs (spleen, thymus, cervical lymph nodes) were meshed through 70 µm filters (Corning, Cambridge, MA, USA, 352350), bone marrow was harvested by flushing both femurs and tibiae using a 23g needle with staining buffer, consisting of PBS with 2% FBS (Gibco, Grand Island, NY, USA, 10270-106) and 10 µg/mL Gentamycin (Gibco 15750-037). Peritoneal lavage was performed post-mortem by injecting 10 mL staining buffer in the peritoneal cavity using a 27g needle, gently massaging the peritoneum and extracting the fluid again with a 27G needle. Erythrocyte depletion was performed by incubating cells for 3 min in 1 mL lysis buffer (155 mM NH4Cl, 10 mM KHCO3, 0.1 mM EDTA; pH7.5) on ice. Cells were washed with staining buffer and filtered through a 50 µm cup falcon (BD Biosciences, San Diego, CA, USA, 340632). Cell numbers were determined using a hemocytometer and trypan blue exclusion.

### Cell sorting

Single-cell suspensions were pre-incubated with 1 µg/mL of aCD16/32 Fc-Block (BioLegend, San Diego, CA, USA, 101310) in 300 µL staining buffer for 10 min, washed and stained for 15 min with antibodies in a volume of 300 µL staining buffer. The sorted cell subsets were defined as follows: **Bone Marrow:** pro B cells (B220loCD19+IgD-IgM-CD25-cKit+), large pre-B cells (B220loCD19+IgD-IgM-CD25+ckit-FSChi), small pre-B cells(B220loCD19+IgD-IgM-CD25+ckit-FSClo), immature IgM+ B cells (B220loCD19+IgD-IgM+), CLP (Lin-cKit^int^Sca1+CD127^hi^) and LSK (Lin-cKit+Sca1+). **Spleen:** Follicular B cells (B220+CD19+AA4.1-IgM+CD21+CD23+), marginal zone B cells (B220+CD19+AA4.1-IgM+CD21+CD23-), transitional B cells (CD19+B220+AA4.1+), CD4/8 naïve (CD3+CD4+/CD8+CD62L+CD44-), CD4/8 CM (CD3+CD4+/CD8+CD62L-CD44+), CD4/8 EM (CD3+CD4+/CD8+CD62L+CD44+). **Thymus:** DP (CD4+CD8+LIN-), DN (CD4-CD8-LIN-).

### Peritoneal Cavity

B1 B cells (B220loCD19+CD43+). Cell sorting was carried out on a FACS Aria III (BD Biosciences). Non-singlet events were excluded from analyses based on characteristics of FSC-H/FSC-W and SSC-H/SSC-W.

### Flow Cytometry

For flow cytometric analysis of lymphatic organs 1 × 10^6^ cells in single-cell suspension were used for each staining. For the analysis of cultured cells, a minimum of 50.000 cells was used. The staining procedure was performed in 96-well plates, and cells were washed with 200 µL staining buffer by centrifugation at 2.000rpm for 2 min. Staining was conducted on ice or at 4°C. To block non-specific antibody-binding, cells were pre-incubated with 1 μg/mL of αCD16/32 Fc-Block (BioLegend 101310) in 30 μl staining buffer (PBS, 2% FCS, 10 µg/mL Gentamycin) for 10 min, washed, and stained for 20 min with 30 μl antibody cocktails. The following fluorescent-labeled anti-mouse antibodies were used at dilution 1:100 – 1:1000

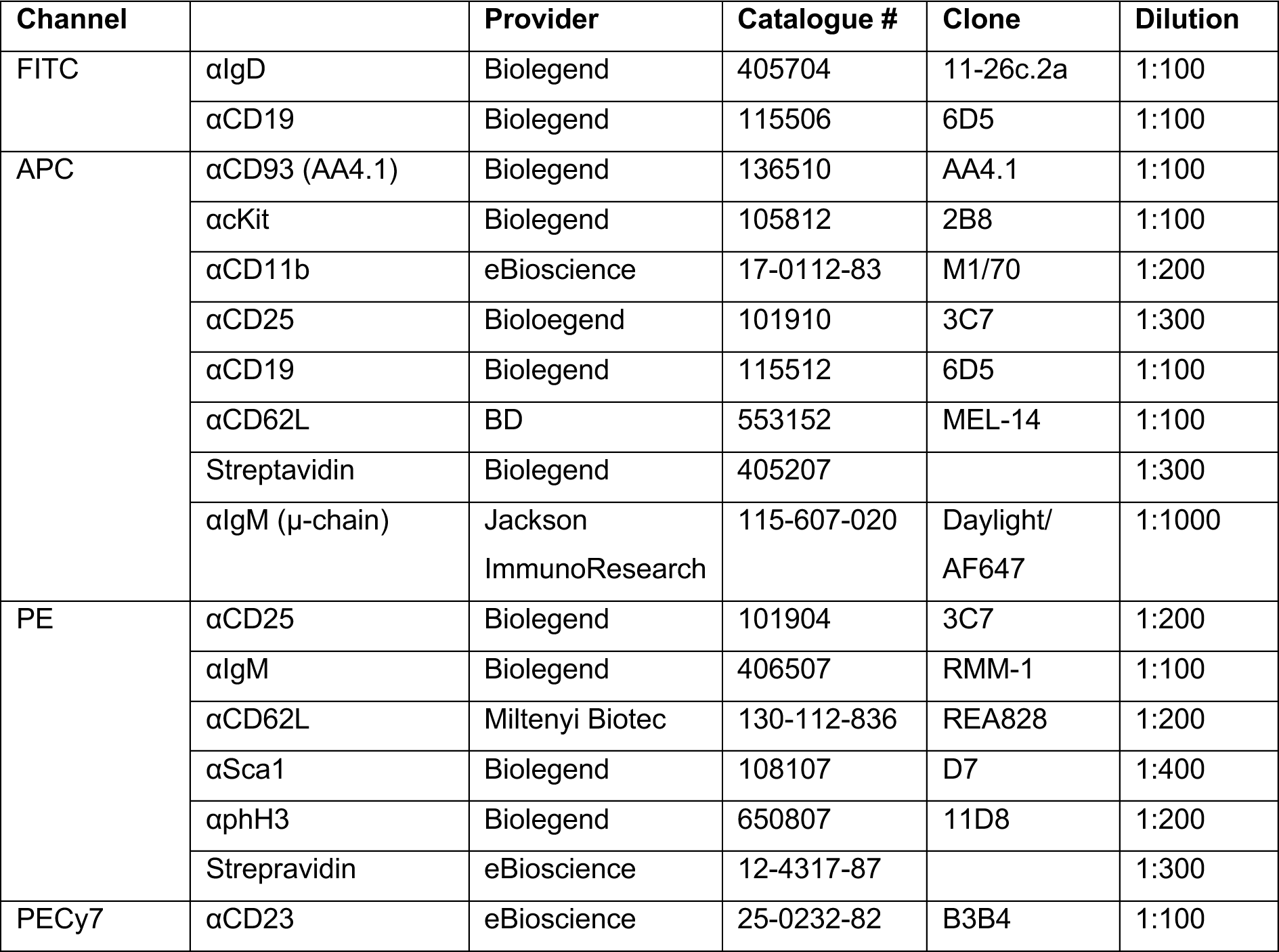
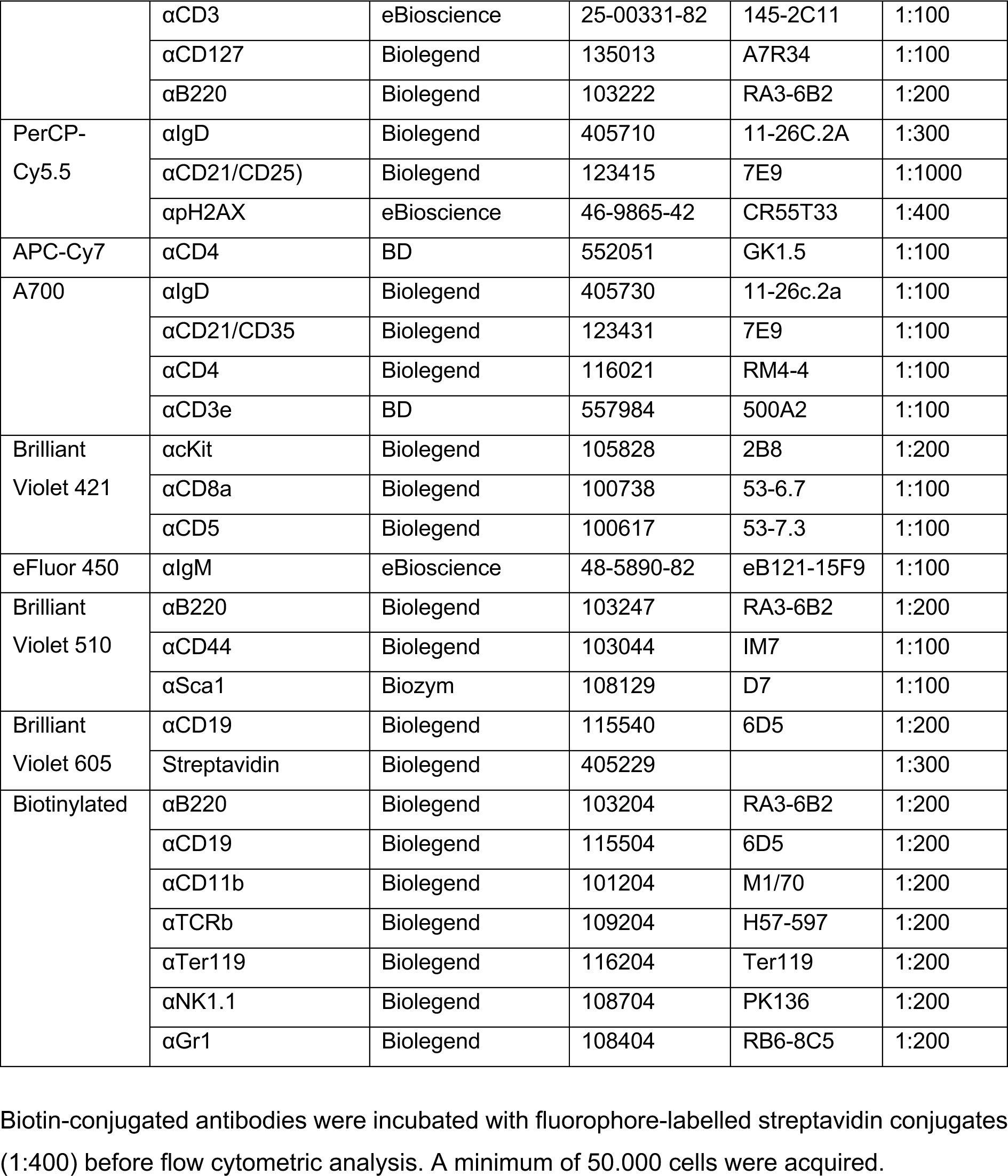

| Channel |  | Provider | Catalogue # | Clone | Dilution |
| --- | --- | --- | --- | --- | --- |
| FITC | αIgD | Biolegend | 405704 | 11-26c.2a | 1:100 |
| | αCD19 | Biolegend | 115506 | 6D5 | 1:100 |
| APC | αCD93 (AA4.1) | Biolegend | 136510 | AA4.1 | 1:100 |
| | αcKit | Biolegend | 105812 | 2B8 | 1:100 |
| | αCD11b | eBioscience | 17-0112-83 | M1/70 | 1:200 |
| | αCD25 | Biolegend | 101910 | 3C7 | 1:300 |
| | αCD19 | Biolegend | 115512 | 6D5 | 1:100 |
| | αCD62L | BD | 553152 | MEL-14 | 1:100 |
|  | Streptavidin | Biolegend | 405207 |  | 1:300 |
| | αIgM (μ-chain) | Jackson ImmunoResearch | 115-607-020 | Daylight/AF647 | 1:1000 |
| PE | αCD25 | Biolegend | 101904 | 3C7 | 1:200 |
| | αIgM | Biolegend | 406507 | RMM-1 | 1:100 |
| | αCD62L | Miltenyi Biotec | 130-112-836 | REA828 | 1:200 |
| | αSca1 | Biolegend | 108107 | D7 | 1:400 |
| | αphH3 | Biolegend | 650807 | 11D8 | 1:200 |
|  | Streptavidin | eBioscience | 12-4317-87 |  | 1:300 |
| PECy7 | αCD23 | eBioscience | 25-0232-82 | B3B4 | 1:100 |
| | αCD3 | eBioscience | 25-00331-82 | 145-2C11 | 1:100 |
| | αCD127 | Biolegend | 135013 | A7R34 | 1:100 |
| | αB220 | Biolegend | 103222 | RA3-6B2 | 1:200 |
| PerCP-Cy5.5 | αIgD | Biolegend | 405710 | 11-26C.2A | 1:300 |
| | αCD21/CD25) | Biolegend | 123415 | 7E9 | 1:1000 |
| | αpH2AX | eBioscience | 46-9865-42 | CR55T33 | 1:400 |
| APC-Cy7 | αCD4 | BD | 552051 | GK1.5 | 1:100 |
| A700 | αIgD | Biolegend | 405730 | 11-26c.2a | 1:100 |
| | αCD21/CD35 | Biolegend | 123431 | 7E9 | 1:100 |
| | αCD4 | Biolegend | 116021 | RM4-4 | 1:100 |
| | αCD3e | BD | 557984 | 500A2 | 1:100 |
| Brilliant Violet 421 | αcKit | Biolegend | 105828 | 2B8 | 1:200 |
| | αCD8a | Biolegend | 100738 | 53-6.7 | 1:100 |
| | αCD5 | Biolegend | 100617 | 53-7.3 | 1:100 |
| eFluor 450 | αIgM | eBioscience | 48-5890-82 | eB121-15F9 | 1:100 |
| Brilliant Violet 510 | αB220 | Biolegend | 103247 | RA3-6B2 | 1:200 |
| | αCD44 | Biolegend | 103044 | IM7 | 1:100 |
| | αSca1 | Biozym | 108129 | D7 | 1:100 |
| Brilliant Violet 605 | αCD19 | Biolegend | 115540 | 6D5 | 1:200 |
|  | Streptavidin | Biolegend | 405229 |  | 1:300 |
| Biotinylated | αB220 | Biolegend | 103204 | RA3-6B2 | 1:200 |
| | αCD19 | Biolegend | 115504 | 6D5 | 1:200 |
| | αCD11b | Biolegend | 101204 | M1/70 | 1:200 |
| | αTCRb | Biolegend | 109204 | H57-597 | 1:200 |
| | αTer119 | Biolegend | 116204 | Ter119 | 1:200 |
| | αNK1.1 | Biolegend | 108704 | PK136 | 1:200 |
| | αGr1 | Biolegend | 108404 | RB6-8C5 | 1:200 |

Biotin-conjugated antibodies were incubated with fluorophore-labelled streptavidin conjugates (1:400) before flow cytometric analysis. A minimum of 50.000 cells were acquired.

### Intracellular staining and DNA content analysis

Fixation was performed in 200 µL ice-cold 70% ethanol for a minimum of 16 hours at −20 °C in 96-well plates. Prior antibody staining, cells were washed twice adding 200 µL PBS (2000 r.p.m., 5 min) to remove ethanol, transferred, and subsequently permeabilized with 200 μl PBS containing 0.25% Triton X-100 (Sigma-Aldrich, St. Louis, MO, USA, T8787) for 20 min. Cells were incubated with ɣH2AX (Cell Signaling, Beverley, MA, USA, 2577, 1:400) and phospho-Histone H3 Ser-10 (Cell Signaling 9701, 1:400) antibody in 30 μl permeabilization buffer for 20–30 min. Cells were washed twice with permeabilization buffer, resuspended in 100 μl PBS containing 250 μg/mL RNase A (Sigma R5500), and incubated for 20 min at 37 °C. Finally, 50 μl of 3 μM TO-PRO-3 or 50 μl 10 μg/mL DAPI (Sigma D9542) in PBS was added. A minimum of 10.000 cells was acquired at LOW mode. Data were acquired on an LSRII cytometer (BD Biosciences) and analysed using FLOWJO software (Tree Star, Ashland, OR, USA). Non-singlet events were excluded from analyses using FSC-H/FSC-W and SSC-H/SSC-W characteristics and in case of DNA staining in addition with DAPI 405-A/405-H or TO-PRO-3 APC-A/APC-H.

### Pro B cell culture

To expand pro-B cells *ex vivo*, 2-4 x 10^5^ B220^lo^CD19^+^IgD^-^IgM^-^CD25^-^ cKit^+^cells were FACS-sorted from the bone marrow of approximately 5-week-old mice. One third of the cells was used for cell cycle analysis and centriole count, the remaining cells were then plated in in 96-well plates in B cell Medium (RPMI-1640 medium (Thermo Fisher Scientific, Waltham, MA, USA, 21875034) supplemented with 10% FBS, 2 mM L-glutamine (Sigma-Aldrich G7513), 0.055 mmol/L 2-ME (Thermo-Fisher 31350010), 10 mmol/L HEPES (Sigma-Aldrich H0887), 1 mmol/L sodium pyruvate (Gibco, 11360-039), 1 x NEAA (Gibco, 11140-035), 100 units/mL penicillin, and 100 μg/mL streptomycin (Sigma-Aldrich, P0781) together with 20 ng/mL IL-7 (Peprotech 217-17). A separate 96-well was prepared for the analysis of each day/condition. Cells were harvested by pipetting, washed once with PBS and then split equally for cell cycle analysis and microscopy (or qPCR). Doxycyclin (Sigma-Aldrich D9891, 1 µg/mL) was added to the culture medium 48h after plating. Centrinone (LCR-263, MedChemExpress HY-18682) was added directly after plating the cells.

### Induced Germinal Center culture (iGC) B cell culture

B cells were isolated from splenic single-cell suspension using MagniSort Streptavidin Negative Selection Beads (Thermo Fisher Scientific, MSNB-6002-74) and biotinylated antibodies against Ter119 (BioLegend, San Diego, CA, USA, 116204, 1: 100), CD11b (BioLegend, 101204, 1: 100) and TCRβ (BioLegend, 109204, 1: 50). The iGC B cell culture was performed as described by Nojima et al. (Citation: https://pubmed.ncbi.nlm.nih.gov/21897376/). Briefly, 4×10^5^ 40LB feeder cells were treated with 10µg/ml mitomycin C (Sigma, M0305) in 1ml DMEM (Sigma, WHMISDZB) supplemented with 10% FBS (Gibco, 10270-106), 2 mm l-glutamine (Sigma, G7513) and 100 U·mL−1 penicillin/100 μg·mL−1 streptomycin (Sigma, 0781). Cells were washed five times with PBS and then, 7,5×10^5^ B cells were added in B cell medium: DMEM supplemented with 10% (v/v) FBS, 2 mm l-glutamine, 10 mm Hepes (LONZA, Basel, Switzerland, BE17-737E), 1 mm sodium pyruvate (Gibco, 13360-039), 1× nonessential amino acids (Gibco, 11140-035), 100 U·mL−1 penicillin/100 μg·mL−1 streptomycin, 50 μm β-mercapto-ethanol (Sigma, M3148), 10 ng·mL−1 rIL-4 (Peprotech, Rocky Hill, NJ, USA, 214-14) and treated with different concentrations of centrinone. On day 4, iGC B cells were harvested by collecting the medium and washing plates with harvest buffer (PBS with 0.5% BSA and 2mM EDTA). Cells were either analysed or 7.5 × 10^5^ iGC B cells were replated per 6-well containing fresh mitomycin C-treated 40LB feeder cells as detailed above, and cultivated in 4 mL of B-cell medium containing either 10 ng·mL−1 rIL-4 or 10 ng·mL−1 rIL-21 (Peprotech, 210-21). On day 8, B cells were harvested and used for flow cytometric analysis.

For surface flow cytometric analysis iGC B cells were stained as described in flow cytometry section with the following antibodies (B220 PE 1:1000, IgM PECy7 1:200, IgD PerCPCy5.5 1:400, CD22 APC 1:200, IgG1 A700 1:200, CD19 BV605 1:200, CD138 BV510 1:200). Before acquisition cells were labelled with Fixable Viability Dye eFluor 780 (Thermo Fisher Scientific, 65-0865-14) as per manufacturer’s instructions. For intracellular flow cytometric analysis, iGC B cells were washed once with PBS and then treated with trypsin for 10min at 37°C. Subsequently, cells were washed with staining buffer and fixed by the addition of self-made fixation solution (PBS + 4%PFA + 0,1% Saponin) for 20min at 4°C. Cells were washed two times with Perm/Wash (PBS + 1% BSA +0,1% Saponin + 0,025% NatriumAzid) and then incubated with the following antibodies for 15min (IgG1 FITC 1:100, pH3 PE 1:400, IgM PECy7 1:200, gH2AX PerCP Cy5.5 1:400, IgE Bio 1:200). After washing with perm/wash buffer cells were incubated with second antibody solution (Strep BV605 1:400) and after 15min incubation further processed as described in the section intracellular staining and DNA content analysis.

### Immunofluorescence microscopy

For the imaging of B cells from peripheral blood, blood was collected from the submandibular vein into EDTA coated tubes (BD 365974) and red blood cells were lysed in RBC Lysis Buffer (eBioscience^TM^ 00-4333-57) for 5 min at room temperature. Tubes were centrifuged at 600 x g for 5 min and pellets were washed in PBS 2 % FBS, before incubation with CD19 Alexa Fluor 647 (BioLegend 115522, 1:100) for 30 min at room temperature.

In general, cells were washed once with PBS and transferred to Poly-L-Lysine (Sigma-Aldrich P8920) coated coverslips, followed by incubation for 30 min at 4°C or poly-D-lysine (Sigma-Aldrich P6407) coated coverslips, followed by incubation at 37°C for 1 hour. Cells were then fixed with 100 % ice-cold methanol for 10 min at −20°C and washed 3 times with PBS. Cells were stored for up to two weeks in PBS or stained immediately. Cells were incubated with blocking solution (2.5 % FBS, 200 mM glycine, 0.1 % Triton X-100 in PBS or 2 % bovine serum albumin (GE Healthcare or Calbiochem), 0.01 % Tween 20 (National Diagnostics or Roth) in PBS) for 1 h at room temperature before staining with primary antibodies in blocking solution for 1 hour. Cells were washed 3 x with washing buffer (PBS 0.1 % Triton X-100 or PBS with 0.2 % Tween) and then incubated with secondary antibodies. When DAPI was used, it was added 1:1000 in secondary antibody dilution. Cells were again washed 3 x with washing buffer. When Höchst 3342 (Sigma-Aldrich 23491-52) was used it was added in the last washing step. Coverslips were mounted in ProLong^TM^ Gold Antifade (ThermoFisher Scientific) or home-made mounting solution (23 % polyvinyl alcohol, 63 % glycerol and 0,02 % NaN3 in PBS). The following antibodies were used to perform immunofluorescence staining in murine cells: rabbit polyclonal α-CEP152 1:1000 (homemade, LoMastro *et al*., 2022, 1:1000), goat polyclonal α-ɣ-Tubulin (homemade, Levine *et al*., 2017, 1:1000), rabbit polyclonal α-CEP135 (homemade, LoMastro *et al*., 2022, 1:1000), mouse α-ɣ-Tubulin (Sigma-Aldrich, 1:250); rabbit α-CP110 (Protein Tech, 12780-1-AP, 1:500), goat α-mouse IgG AF568 (Thermo Fisher, A11031, 1:1000), goat α-rabbit IgG AF488 (Thermo Fisher, A-11034, 1:1000).

Thirty to fifty Z-stacks were acquired at room temperature on Zeiss Axiovert 200M microscope with an oil immersion objective (Ph3 Plan-Neofluar 100×/1.3 oil, 440481, Zeiss) using the acquisition software VisiView 4.1.0.3. Maximum stack projections of z-stacks were performed using the VisiView 4.1.0.3. ImageJ was used to adjust contrast and brightness and centriole numbers were evaluated manually. CP110+ spots per cell were only counted, if co-localization with ɣ-Tubulin was given. A minimum of 150 cells was counted for each condition.

### Ultrastructure expansion microscopy (U-ExM)

B cells were isolated from mouse spleens using EasySep Mouse B Cell Isolation Kit (Stem Cell Technologies 19854) and attached to poly-D-lysine coated coverslips at 37°C for 1 hour. U-ExM was carried out as previously described (Gambarotto et al., 2021). Coverslips with unfixed cells were incubated in a solution of 0.7 % formaldehyde with 1 % acrylamide in PBS overnight at 37°C. Gelation was carried out via incubation of coverslips with cells facing down with 35 µL of U-ExM MS composed of 19 % (wt/wt) sodium acrylate, 10 % (wt/wt) acrylamide, 0.1 % (wt/wt) N,N′-methylenbisacrylamide (BIS) in PBS supplemented with 0.5 % APS and 0.5 % TEMED, on Parafilm in a pre-cooled humid chamber. Gelation proceeded for 10 min on ice, and then samples were incubated at 37°C in the dark for 1 hour. A 4 mm biopsy puncher (Integra; 33–34 P/25) was used to create one punch per coverslip. Punches were transferred into 1.5 mL Eppendorf tubes containing 1 mL denaturation buffer (200 mM SDS, 200 mM NaCl, and 50 mM Tris in ultrapure water, pH 9) and incubated at 95°C for 1 hr. After denaturation, gels were placed in beakers filled with ddH2O for the first expansion. Water was exchanged at least two times every 30 min at RT, and then gels were incubated overnight in ddH2O. Next, to remove excess water before incubation with primary antibody solution, gels were placed in PBS two times for 15 min. Primary antibodies were diluted in 2 % PBS/BSA and incubated with gels at 37°C for 2.5 hours, with gentle shaking. Gels were then washed in PBST three times for 10 min with shaking and subsequently incubated with secondary antibody solution plus DAPI diluted in 2 % PBS/BSA for 2.5 hour at 37°C with gentle shaking. The gels were then washed in PBST three times for 10 min with shaking and finally placed in beakers filled with ddH2O for expansion. Water was exchanged at least two times every 30 min, and then gels were incubated in ddH2O overnight. Gels expanded between 4.0× and 4.5× according to SA purity. The following antibodies were used for expansion microscopy: mouse monoclonal α-acetylated-α-Tubulin (Cell Signaling Technology 12152, 1:500), rabbit polyclonal α-Centrin (homemade, Moyer et al., 2019, 1:500), Alexa fluor secondaries 1:800 (XXX). And DAPI (1:800) was used to image DNA.

### Quantitative real-time-PCR

Total RNA was extracted from snap-frozen cell pellets using the Quick-RNA Micro Prep Kit (Zymo Research, Irvine, CA, USA, R1050) and DNase digestion as per manufacturer’s instructions. RNA concentrations were measured by Nanodrop and 100ng of total RNA where subjected to first-strand cDNA synthesis with iScript cDNA Synthesis Kit (Bio-Rad, Hercules, CA, USA, 170-8891). cDNA was amplified using the Luna Universal qPCR Master Mix (New England Biolabs M3003E) as per manufacturer’s instructions. The qRT-PCR was run on a StepOnePlus Real-time PCR system (Applied Biosystems, Foster City, CA, USA). Gene expression of individual mRNAs was normalized to HPRT using the ΔC(t) method. The following primers were used:

*Hprt* F: 5’-GTCATGCCGACCCGCAGTC-3’,

*Hprt* R: 5’-AGTCCATGAGGAATAAAC-3’,

*Plk4* F: 5’-GGAGAGGATCGAGGACTTTAAGG-3’

*Plk4* R: 5’-CCAGTGTGTATGGACTCAGCTC-3’

*Plk2* F: 5’-CTGAAGGTGGGAGACTTTG-3’

*Plk2* R: 5’-AGGACTTCGGGGGAGAGATA-3’

*Plk1* F: 5’-CCCTATTACCTGCCTCACCA-3’,

*Plk1* R: 5’-ACCACCGGTTCCTCTTTCTC-3’;

### ELISA

For total immunoglobulin levels in serum 96-well enzyme-linked immunosorbent assay plates (Sigma, CLS3590) were coated with 50 µg/mL capture antibody (Southern Biotech, Birmingham, AL, USA, 1010-01) at 4°C over night. Plates were washed three times with wash buffer (PBS containing 0.05% TWEEN20), blocked with 100 µL per well 1% BSA in PBS at room temperature for 4h and washed three more times with wash buffer. 100 µL per well of mouse serum serially diluted 1:4 in blocking buffer (range 1:800 to 1:160 000) were incubated with coated wells over night at 4°C with. Plates were washed three times with wash buffer and incubated with 100 µL per well of HRP-conjugated α-mouse IgG1 (Southern Biotech, 1070-05, 1:5000 in 1 % BSA in PBS) or HRP-conjugated α-mouse IgM (Southern Biotech, 1020-05, 1:5000 in 1 % BSA in PBS) for 4 h at room temperature. For detection, 100 µL of ABTS substrate solution per well [200μL ABTS (Stock: 15 mg/mL in a.d.), 10 mL citrate-phosphate buffer (574 mg citric acid monohydrate in 50 mL a.d.) and 10 µL H2O2] was incubated for 20 min. Absorbance was measured at 405 nm using a microplate reader (Tecan Sunrise, Männedorf, Switzerland). For OVA-specific serum IgG1 titers 13 days after immunization with OVA/alum, Cayman’s Anti-Ovalbumin IgG1 (mouse) ELISA Kit (Cayman Chemical, Ann Arbor, MI, USA, 500830) was used according to manufacturer’s protocol.

### Quantification and statistical analysis

Results are always shown as mean and standard deviation (SD). Graphs were plotted and statistical analysis was performed with GRAPHPAD PRISM 9.3.1 software (GraphPad Software, San Diego, CA, USA) using unpaired Student’s t test when comparing two groups, One-way ANOVA and Tukey post hoc test when comparing multiple groups and Two-way ANOVA when comparing multiple groups over different timepoints or conditions. All relevant comparisons were made, and non-significant results were not indicated in the figures. The number of biological repetitions (n) is stated in each figure legend, and every experiment was performed at least twice. Differences between groups were considered statistically significant when P < 0.05. In figures, asterisks stand for: *P < 0.05; **P < 0.01; ***P < 0.001; ****P < 0.0001

## Acknowledgements

We thank C. Soratroi, I. Gaggl and J. Heppke for expert technical assistance, S Gritsch and M. Sauerwein for animal care, as well as S. Geley, for help with microscopy and critical discussion. This work was supported by the ERC-AdG “POLICE” (#787171) to AV. MS acknowledges support by the CBD Doctoral College, funded by the Austrian Science Fund, FWF (DOC82).

## Author contributions

MS conducted experiments, analysed data, prepared figures, wrote manuscript. VB, GLM, MSL, conducted experiments and managed mouse colonies. MH provided crucial reagents, AJH, VL & AV designed research, analysed data, wrote manuscript, conceived study.

