## Supplementary Figure 1-6 for "Exact centriole counts are critical for B cell development but not function"

### Supplementary Material

#### Figure S1

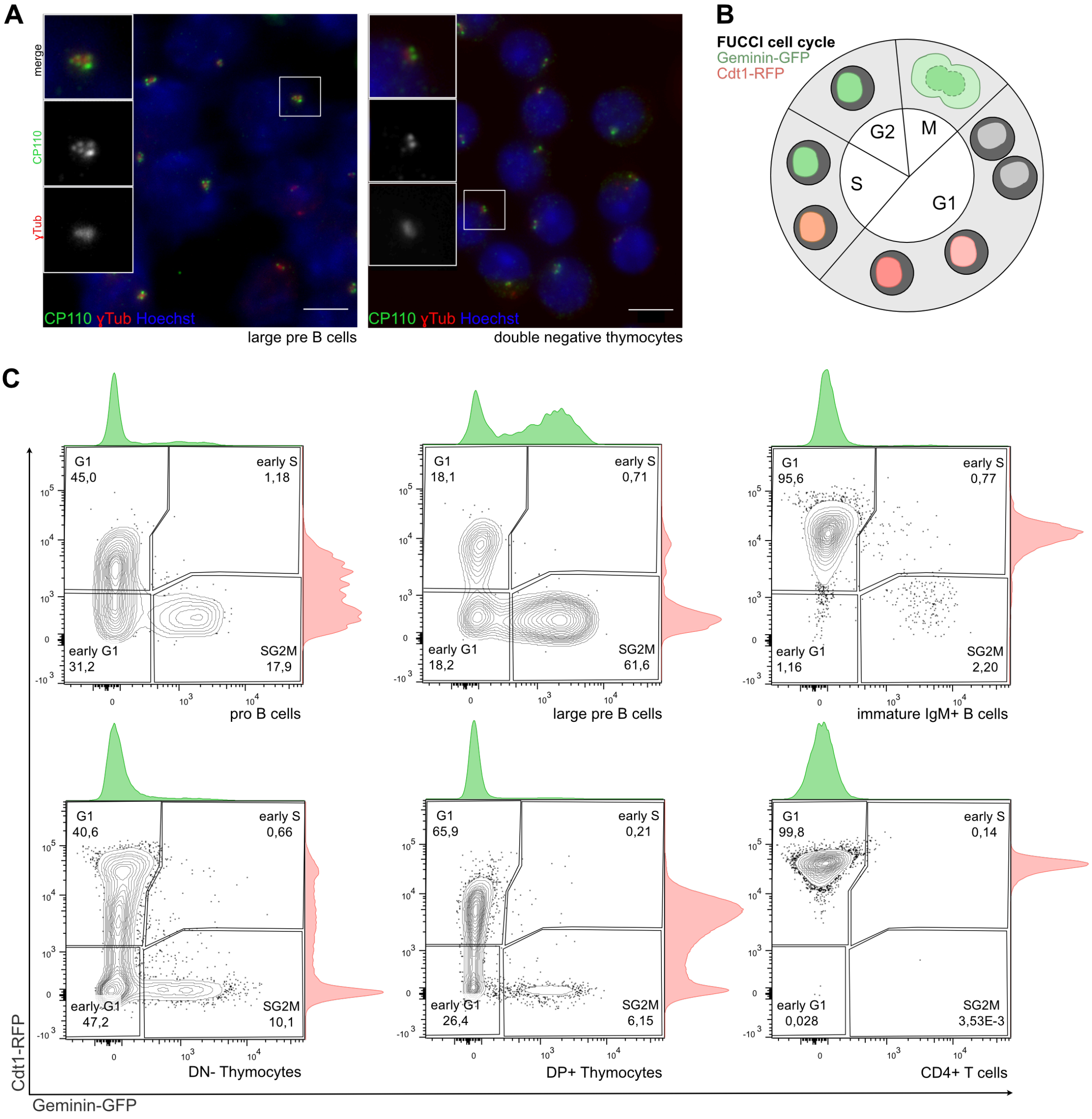


**Fig. S1 | Centriole count correlates with proliferative potential in B progenitor cells**

A. Immunofluorescence images of large pre B cells and double negative thymocytes FACS-sorted from bone marrow and thymus of wild type mice. Cells were stained with CP110, ɣ-Tubulin antibodies and Hoechst. Scale bar, 5µm.

B. Schematic representation of nuclei labeling in cells isolated from FUCCI mice.

C. Representative flow cytometry plots depicting the cell cycle profile of FUCCI mice expressing Geminin-GFP and Cdt1-RFP. Respective cell types are indicated at the bottom of each plot. Bone Marrow: pro B, large and immature IgM+ B-cells. Thymus: double-negative (DN-) and double positive (DP+) thymocytes. Spleen: CD4+ T-cells.

#### Figure S2

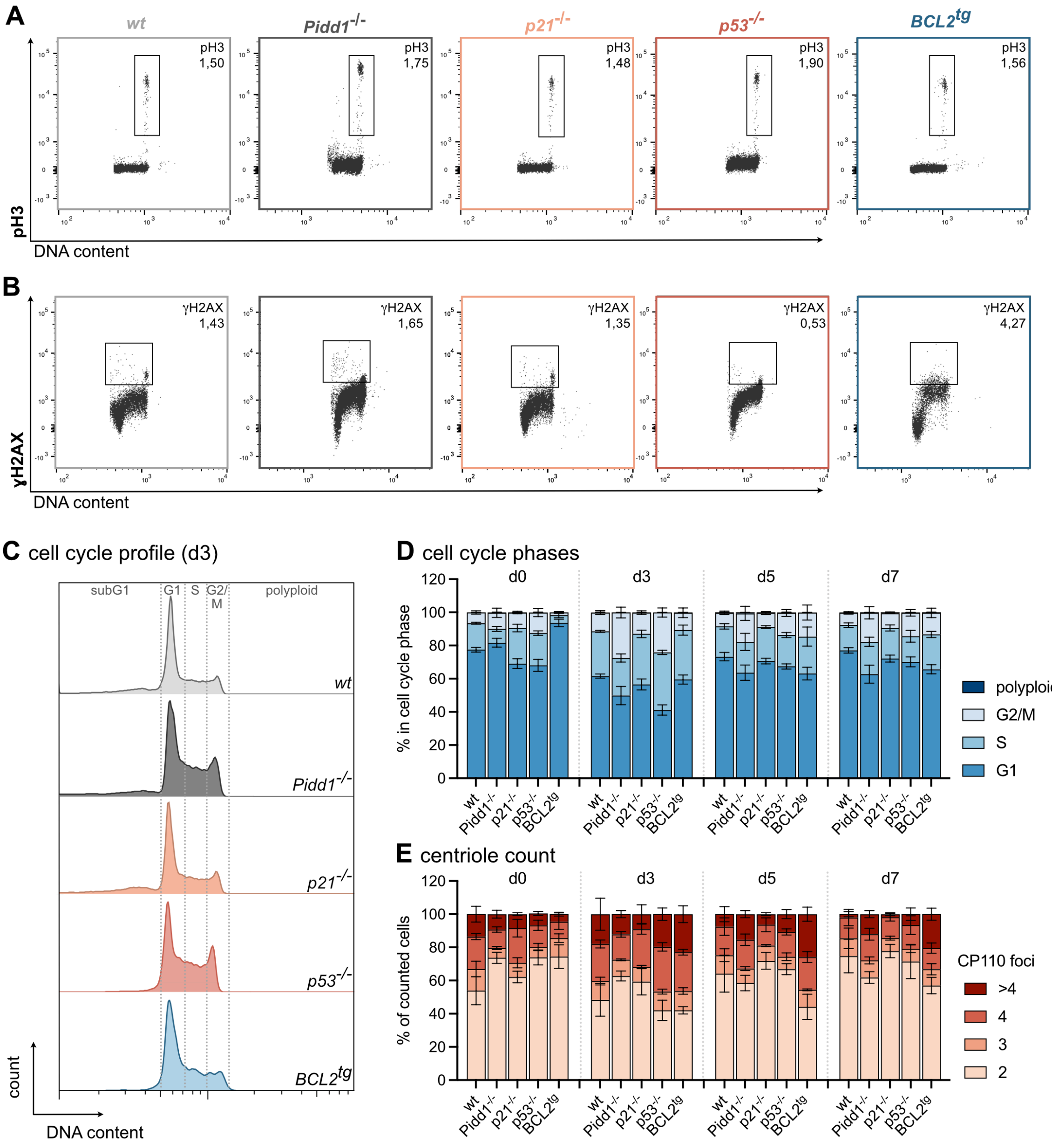


**Fig. S2 | Cell cycle profiling and centriole counts in progenitor B cell cultures**

A. Representative flow cytometry plots of mitotic pro B cells and (B) pro B cells with DNA double strand breaks after 3 days in culture with IL-7 of the indicated genotypes.

C. Flow cytometric histograms depicting the cell cycle profiles of FACS-sorted pro B cells after 3 days in culture with IL-7 of indicated genotypes.

D. Cell cycle profile was determined via flow cytometry directly after sorting (d0) or in culture after 3, 5 and 7 days (d3, d5, d7). Fraction of cells in G1-, S-, G2/M-phase and fraction of polyploid cells are depicted.

E. Percentage of cells with 2,3,4 or more than 4 CP110 foci of pro B cells, determined by immunoflourescence with ɣ-Tubulin and CP110 antibody staining and Hoechst nuclear staining.

Data are shown as mean ± SD; *wt* (n=4-6), *Pidd1*$^{-/-}$ (n=4), *p21*$^{-/-}$ (n=3), *p53*$^{-/-}$ (n=5), *BCL2*$^{tg}$ (n=4).

### Figure S3

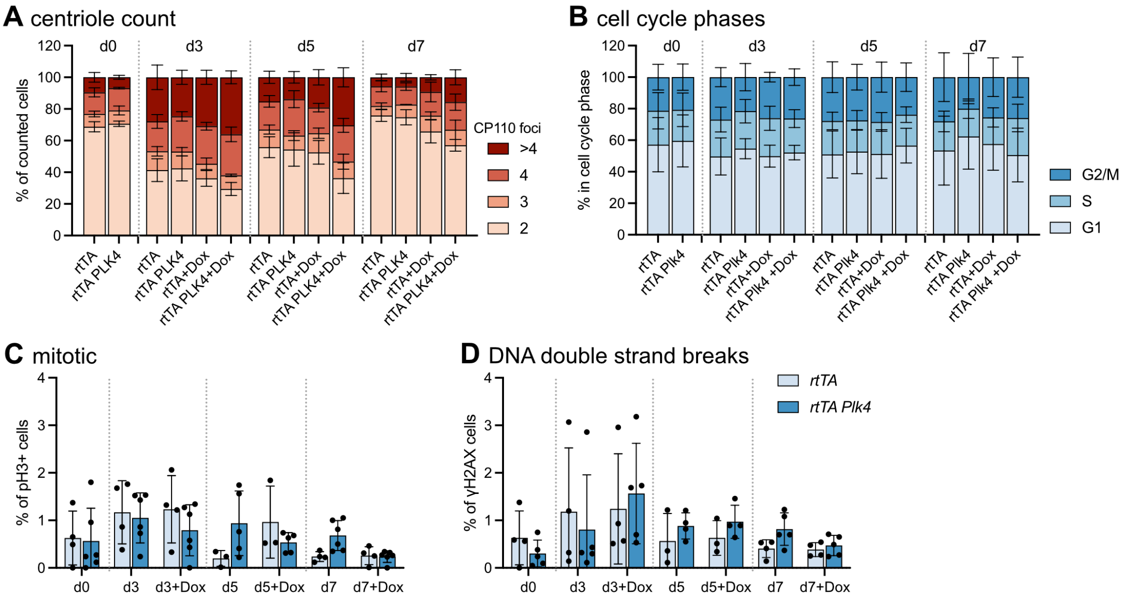


**Fig. S3 | Cell cycle profiling and centriole counts in progenitor B cells overexpressing PLK4**

A. Pro B cells were FACS-sorted from the indicated genotypes and put in culture. Doxycyclin was added after 48h in culture. Fraction of counted pro B cells with 2,3,4 or more than 4 CP110 foci were determined by immunofluorescence with ɣ-Tubulin, CP110 antibodies and Hoechst staining (n=3-6).

B. Fraction of cells in G1-, S-, G2/M-phase of the cell cycle was determined by flow cytometric cell cycle analysis. (n=6 *rtTA*, n=8 *rtTA Plk4* ).

C-D. Fraction of cells positive for (C) mitotic marker pH3 or (D) DNA damage marker ɣH2AX determined by flow cytometric analysis (n=4-6).

Data are shown as mean ± SD; *p<0.05, **p<0.01, ***p<0.001, ****p<0.0001; Two-way-ANOVA Tukey's multiple comparisons test.

#### Figure S4

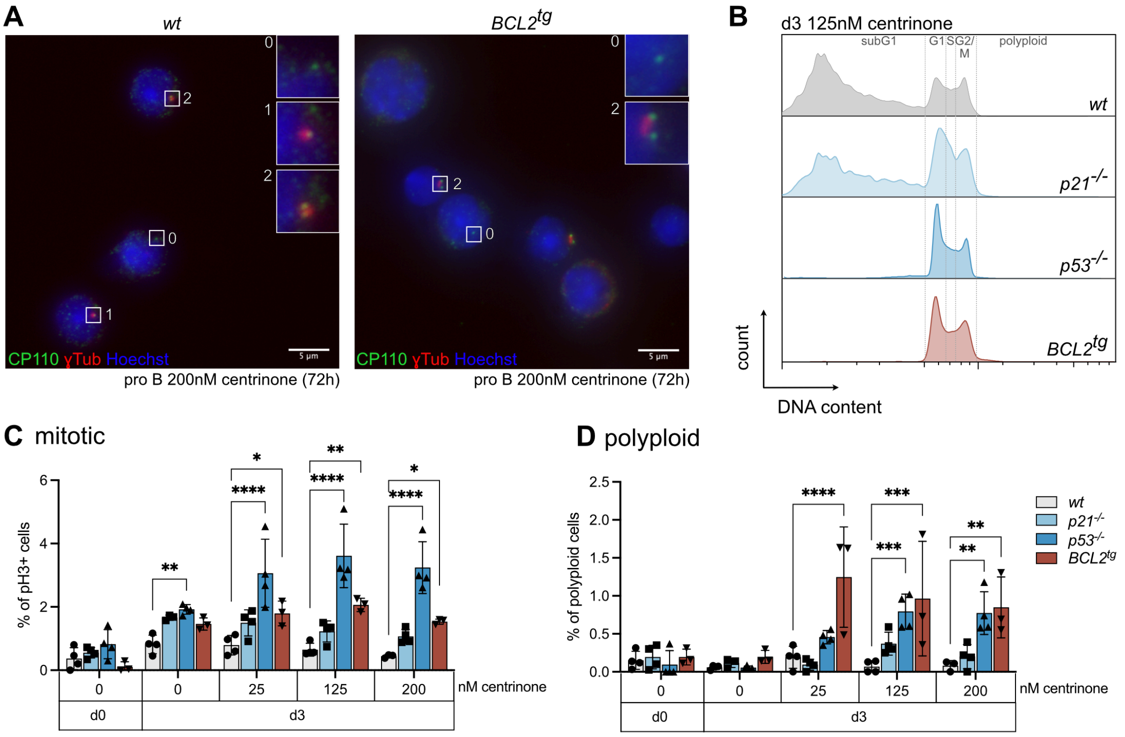


**Fig. S4 | PLK4 inhibition depletes centrioles in progenitor B cells**

A. Representative immunofluorescence-images of *wt* and *BCL2*$^{tg}$ pro B cells treated for 3 days with 200nM centrinone stained with γ-Tubulin, CP110 antibodies and Hoechst. Scale bar, 5µm.

B. Flow cytometric histograms depicting the cell cycle profiles of FACS-sorted pro B cells after 3 days in culture with IL-7 and 125nM centrinone of the indicated genotypes.

C. Fraction of pro B cells positive for mitotic marker pH3 determined by flow cytometric analysis.

D. Fraction of polyploid cells was assessed via flow cytometric DNA content analysis.

Data are shown as mean ± SD; n=3-5; *p<0.05, **p<0.01, ***p<0.001, ****p<0.0001; Genotypes were compared to *wt* by Two-way-ANOVA Tukey's multiple comparisons test.

### Figure S5

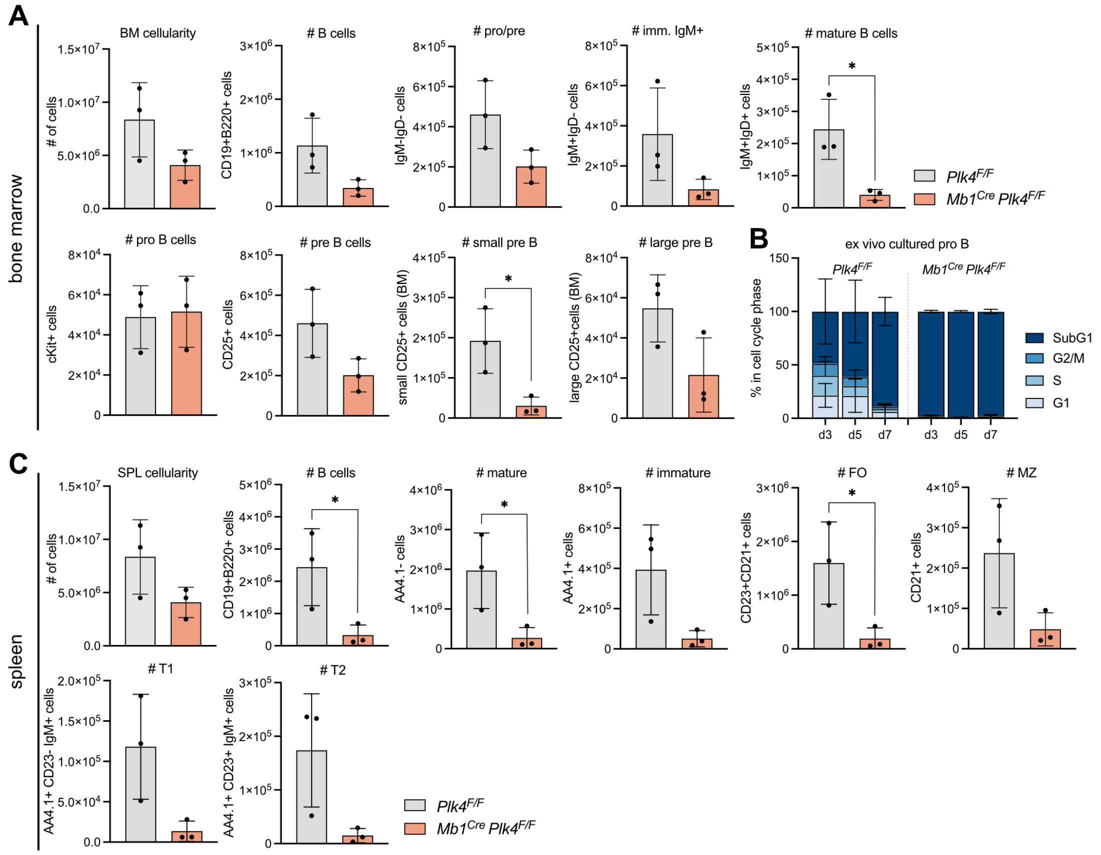


**Fig. S5 | Centrosome loss impairs B cell development**

A. Bone marrow cellularity and total counts of *Plk4$^{F/F}$* or *Mb1$^{Cre}$ Plk4$^{F/F}$* B cells, pro/pre, immature IgM+ and mature B cells, pro B, pre B, small and large pre B cells in the bone marrow.

B. Fraction of FACS-sorted pro B cells in G1-, S-, G2/M-phase of the cell cycle after 3, 5, and 7 days in culture (d3, d5, d7) was determined by flow cytometric cell cycle analysis.

C. Spleen cellularity and total counts of *Plk4$^{F/F}$* or *Mb1$^{Cre}$ Plk4$^{F/F}$* B cells, mature, immature, follicular (FO), marginal zone (MZ), transitional 1 (T1) and 2 (T2) cells in the spleen. Error bars depict the standard deviation of the mean; n=3; *p<0.05, **p<0.01, ***p<0.001, ****p<0.0001; Unpaired T-test.

#### Figure S6

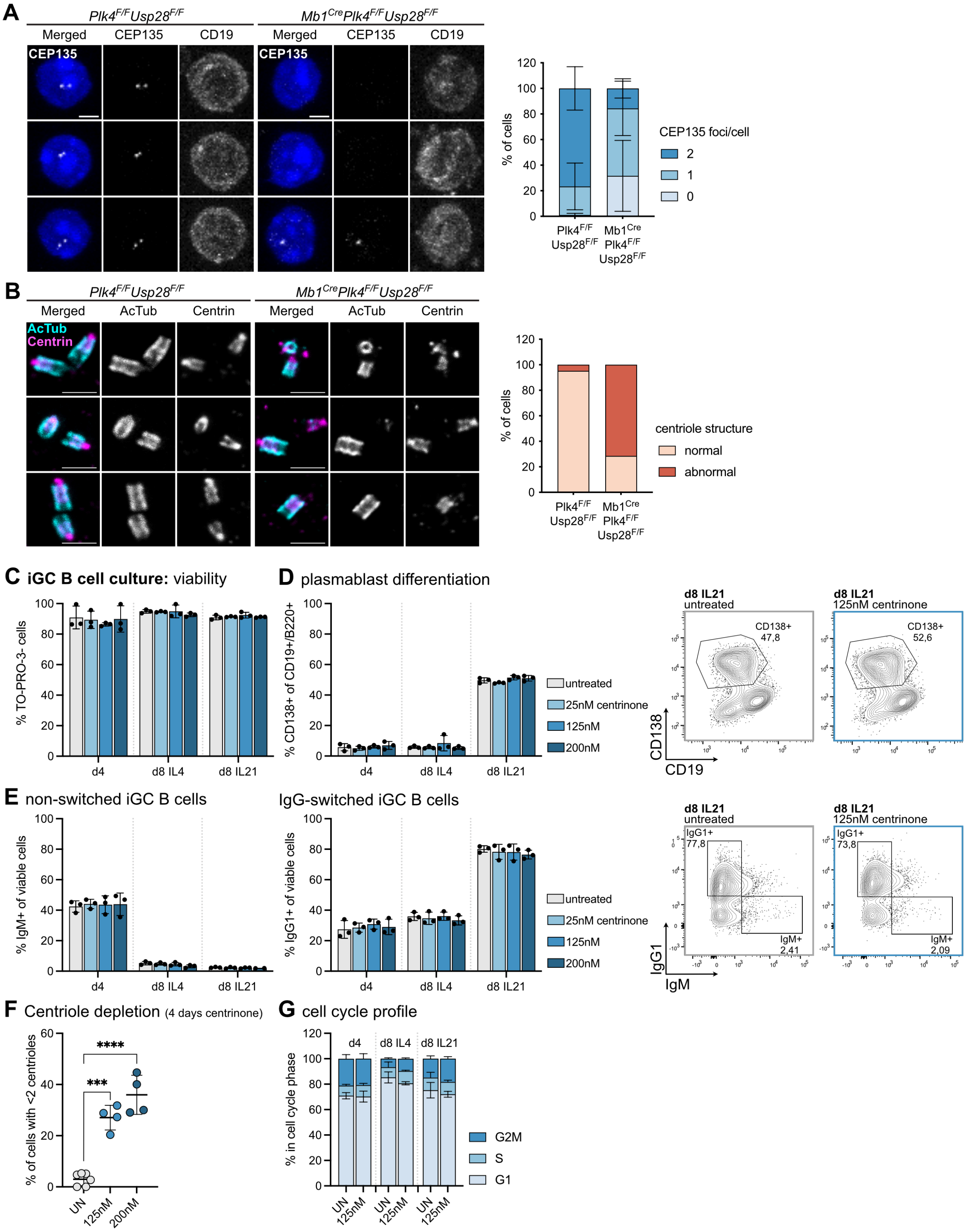


**Fig. S6 | Loss of PLK4 in B cells leads to centriole depletion and structural defects**

A. Number of CEP135 foci per/cell was determined by immunofluorescence of CD19+ B cells of the peripheral blood isolated from *PLK4$^{F/F}$ Usp28$^{F/F}$* and *Mb1$^{Cre}$ Plk4$^{F/F}$ Usp28$^{F/F}$* mice (n=3). Cells were stained with CD19, CEP135 antibodies and DAPI (representative IF images). Scale bars represent 5µm. Data are shown as mean ± SD.

B. Expansion microscopy was used to determine normal and abnormal structure of centrioles in splenic B cells of *PLK4*$^{F/F}$ *Usp28*$^{F/F}$ and *Mb1*$^{Cre}$ *Plk4*$^{F/F}$ *Usp28*$^{F/F}$ mice (n=21 cells of 1 mouse). Cells were expanded by a factor of 4 and stained for acetylated Tubulin (AcTub) and centriole marker Centrin. Scale bars represent 500nm.

C. Flow cytometry analysis of viable (TO-PRO3-negative) B cells cultured on 40LB feeder cells with IL-4 for 4 days, and additional 4 days with either IL-4 or IL-21, treated with graded concentrations of centrinone (25, 125, 200nM).

D. Bar graph (left) and representative flow cytometry plots (right) depicting CD138+ plasmablasts within total iGC B cell cultures (n=3 for each treatment).

E. Bar graphs (left) and representative flow cytometry plots (right) depicting the fraction of IgM+ or switched IgG+ cells within total iGC B cell cultures (n=3 for each treatment).

F. H. Centriole depletion was assessed by immunofluorescence with ɣ-Tubulin, CP110 antibodies and Hoechst staining on iGC B cells after 4 days in culture with 0, 125 or 200nM centrinone (n=6 untreated, n=4 treatment).

G. Intracellular flow cytometry analysis for DNA content performed on untreated and 125nM centrinone treated iGC B cell cultures (n=3 for each treatment).
